# Suppression of cyclooxygenase-2 predisposes to heart failure with preserved ejection fraction

**DOI:** 10.1101/2024.09.28.615616

**Authors:** Emanuela Ricciotti, Philip G. Haines, Manu Beerens, Uri Kartoun, Cecilia Castro, Soon Yew Tang, Soumita Ghosh, Ujjalkumar S. Das, Nicholas F. Lahens, Tao Wang, Jules L. Griffin, Stanley Y. Shaw, Calum A. MacRae, Garret A. FitzGerald

**Author notes:** To whom correspondence should be addressed (E.R.); (G.A.F.). Present address: Rhode Island Hospital, Warren Alpert Medical School of Brown University; Providence, RI. Present address: Center for Computational Health, IBM Research; Cambridge, MA, USA. Present address: Stanford University; Stanford, CA, USA. Present address: One Brave Idea, Division of Cardiovascular Medicine, Brigham and Women’s Hospital; Boston, MA, USA.

## Abstract

Heart failure (HF) is one of the most strongly associated adverse cardiovascular events linked to the use of cyclooxygenase (COX)-2 selective and non-selective nonsteroidal anti-inflammatory drug (NSAID). Nevertheless, it remains uncertain whether NSAID exposure is more likely to lead to heart failure with reduced ejection fraction (HFrEF) or preserved ejection fraction (HFpEF).

In adult mice, postnatal genetic deletion or pharmacological inhibition of COX-2 did not affect cardiac function. In contrast, aged female inducible COX-2 (iCOX-2) knockout (KO) mice displayed diastolic dysfunction, cardiac hypertrophy, pulmonary congestion, and elevated levels of plasma N-terminal pro B-type natriuretic peptide (BNP) when compared to age- and sex-matched controls, while their ejection fraction (EF) remained preserved (≥ 50%). No such phenotype was observed in aged male iCox-2 KO mice. Aged female iCox-2 KO mice showed a shift from prostanoid to leukotriene biosynthesis, along with changes in the expression of mitochondrial genes and calcium-handling proteins in the myocardium. The ratio of phospholamban to SERCA2a was increased, indicating an inhibitory effect on SERCA2a activity, which may contribute to impaired myocardial relaxation. In larval zebrafish, COX-2 inhibition by celecoxib caused a modest yet significant reduction in heart rate and diastolic function, while EF was preserved. Additionally, celecoxib increased BNP expression and ventricular calcium transient amplitude. Diabetic patients in the Harvard-Partners electronic medical record exposed to NSAIDs selective for COX-2 inhibition were more strongly associated with an increased risk of HFpEF compared to HFrEF.

Collectively, these findings indicate that COX-2 deletion or inhibition does not impair systolic cardiac function but instead leads to an HFpEF phenotype in mice, zebrafish, and humans. An imbalance in calcium handling may mediate the impairment of myocardial relaxation following COX-2 suppression.

**Summary:** Genetic deletion or pharmacological inhibition of COX-2 results in heart failure with preserved ejection fraction across zebrafish, mice, and humans.

## Introduction

Nonsteroidal anti-inflammatory drugs (NSAIDs) exert their effects by inhibiting the activity of cyclooxygenase (COX)-1 and COX-2, leading to a reduction of the biosynthesis of prostaglandins (PGs) and thromboxane (Tx). Although they are effective in alleviating pain, inflammation, and fever, their use is constrained by gastrointestinal (GI) and cardiovascular (CV) complications (*1, 2*).

Traditional NSAIDs (tNSAIDs), such as ibuprofen and naproxen, inhibit both COX isozymes. NSAIDs selective for COX-2 inhibition (coxibs), such as rofecoxib and celecoxib, were initially developed to reduce the incidence of serious upper GI complications but have resulted in an increased incidence of adverse CV events (*3, 4*).

A meta-analysis of over 600 randomized clinical trials conducted by the Coxib and traditional NSAID Trialists’ (CNT) Collaboration reported that the use of coxibs and tNSAIDs, except naproxen, was associated with a 30% increased risk of major vascular events (*5*). Moreover, heart failure (HF) was the adverse CV event most strongly linked to both coxibs and tNSAIDs, including naproxen, increasing the risk for hospitalization due to HF by two-fold (*5*). These findings are consistent with epidemiological observations (*6–11*) and randomized controlled trials of coxibs (*12*) and tNSAIDs also selective for COX-2 inhibition (*13*). Additionally, the use of NSAIDs in patients with chronic HF increased the risk of CV adverse events, including a higher risk of recurrent hospitalization due to myocardial infarction (MI) or HF, as well as a dose-dependent increase in the risk of death (*8*).

The increased risk of atherothrombotic events associated with NSAID use is mechanistically consistent with vascular COX-2 inhibition and reduced formation of cardioprotective prostanoids (14). However, the mechanism underlying the association between NSAID exposure and HF risk remains unexplored. The differential impact of naproxen, which causes prolonged and persistent inhibition of platelet COX-1 like aspirin, on atherothrombotic events and HF risk suggests that NSAIDs may contribute to HF through a COX-2-dependent mechanism that is not constrained by variable COX-1-dependent platelet inhibition (*5*).

HF is a heterogeneous clinical syndrome that can be classified by left ventricular EF (*15, 16*). HF is primarily categorized as HF with reduced EF (HFrEF; EF ≤ 40%) or HF preserved EF (HFpEF; EF ≥ 50%). HFrEF, also known as systolic HF, occurs when the left ventricle (LV) loses the ability to contract normally. HFpEF, also known as diastolic HF, occurs when the LV loses the capacity to relax normally. Although HFrEF and HFpEF are equally common, they differ in etiology, pathophysiology, clinical manifestations, and response to therapy (*17*).

The prevalence of HF in the general population increases with age, as does the rate of hospital readmission due to HF (*18, 19*). Elderly patients exposed to NSAIDs exhibit an increased risk of hospitalization (*9, 20*) and death due to HF, as well as recurrent congestive HF (*21, 22*).

Sex-specific differences in cardiac structure, function, and reserve capacity align with the observation that HFrEF is more prevalent in men, while HFpEF is more prevalent in women (*23*). It remains unclear which type of HF is associated with NSAID exposure and whether this relationship is modified by age or sex. Here, we report that COX-2 deletion or inhibition results in diastolic dysfunction in aged female mice and zebrafish. The HFpEF phenotype in mice arises from unbalanced calcium handling, which contributes to impaired myocardial relaxation. In diabetic patients, the use of COX-2 selective NSAIDs, compared to nonselective NSAIDs, increase the likelihood of developing HFpEF rather than HFrEF.

## Results

### Cox-2 deletion or inhibition does not affect cardiac function in adult mice

Genetic and pharmacological manipulations in mice have recapitulated the increased predisposition to thrombosis, hypertension, and atherosclerosis consequent to the suppression of COX-2 dependent prostanoids by NSAIDs (*14, 25*). Therefore, to determine the role of COX-2 on cardiac function, we performed comprehensive functional and structural phenotyping of the hearts of mice in which COX-2 was deleted or inhibited.

We examined post-natal tamoxifen inducible Cox-2 KO mice (iCox-2 KO; *25*) and C57BL/6 mice fed a diet containing celecoxib (100 mg/Kg) or rofecoxib (50 mg/Kg) (*26*). iCox-2 KO mice were previously generated to bypass the multiple abnormalities (including kidney defects, cardiac fibrosis and premature death) evident in conventional Cox-2 KOs (*27*).

Since HF risk is already evident during the first months of NSAID therapy (*28*), we phenotyped the mice approximately 2 months after COX-2 deletion or inhibition.

In adult female mice, the plasma concentration of celecoxib was 0.35± 0.02 μM (n=8) and 0.39±0.04 μM (n=10) of rofecoxib. The corresponding concentrations in adult male mice were 0.50±0.10 μM (n=8) of celecoxib and 0.48± 0.05 μM (n=10) of rofecoxib. The drug concentrations did not differ statistically between the sexes and were in the therapeutic range (*26*).

There was no appreciable difference in body weight (Fig. S1A) or in parameters of systolic (EF) and diastolic (E/e’ ratio, a surrogate marker of LV diastolic filling pressures) function or structural remodeling between adult female and male iCox-2 KO mice compared to their age- and sex-matched WT controls (Fig. S1 B-C; Table S1 and S2). Additionally, Cox-2 deletion did not affect baseline systolic blood pressure (SBP) in adult female and male mice as measured by telemetry (Fig. S1D), consistent with previous reports using a tail-cuff system (*25*).

Similarly, we did not observe an effect of celecoxib or rofecoxib on body weight (Fig. S1E) or on parameters of cardiac function and structural remodeling in adult female and male mice compared to age- and sex- matched mice on a control diet (Fig. S1, F-G, Table S3 and S4). SBP was not changed by the two coxibs in adult female and male mice (Fig. S1H), consistent with previous report (*26*).

Collectively, these results indicate that in adult mice, the suppression of COX-2 alone, in the absence of other perturbations, is not sufficient to impair cardiac function or increase blood pressure.

### Cox-2 deletion impairs diastolic function in aged female mice

Since aging is one of the major risk factors for HF (*18, 19*), we characterized cardiac function in approximately 2-year-old female and male iCox-2 KO mice and their age- and sex-matched WT controls.

In aged mice, Cox-2 deletion did not affect body weight (Fig. 1A). Aged iCox-2 KO female mice exhibited preserved EF and a prolonged E/è ratio, consistent with the presence of diastolic dysfunction (Fig. 1, B-C). Other cardiac parameters did not differ between aged WT and iCox-2 KO female mice (Table S5). Increased LV end diastolic pressure (EDP), measured by invasive hemodynamics, both at rest and during a dobutamine stress test, corroborated the phenotype of diastolic dysfunction observed in aged iCox-2 KO female mice (Fig. 1, D-G, Table S6). Both echocardiographic and invasive hemodynamic data failed to reveal cardiac dysfunction in aged iCox-2 KO male mice (Fig. 1, B-G, Table S7 and S8).

**Fig. 1.**
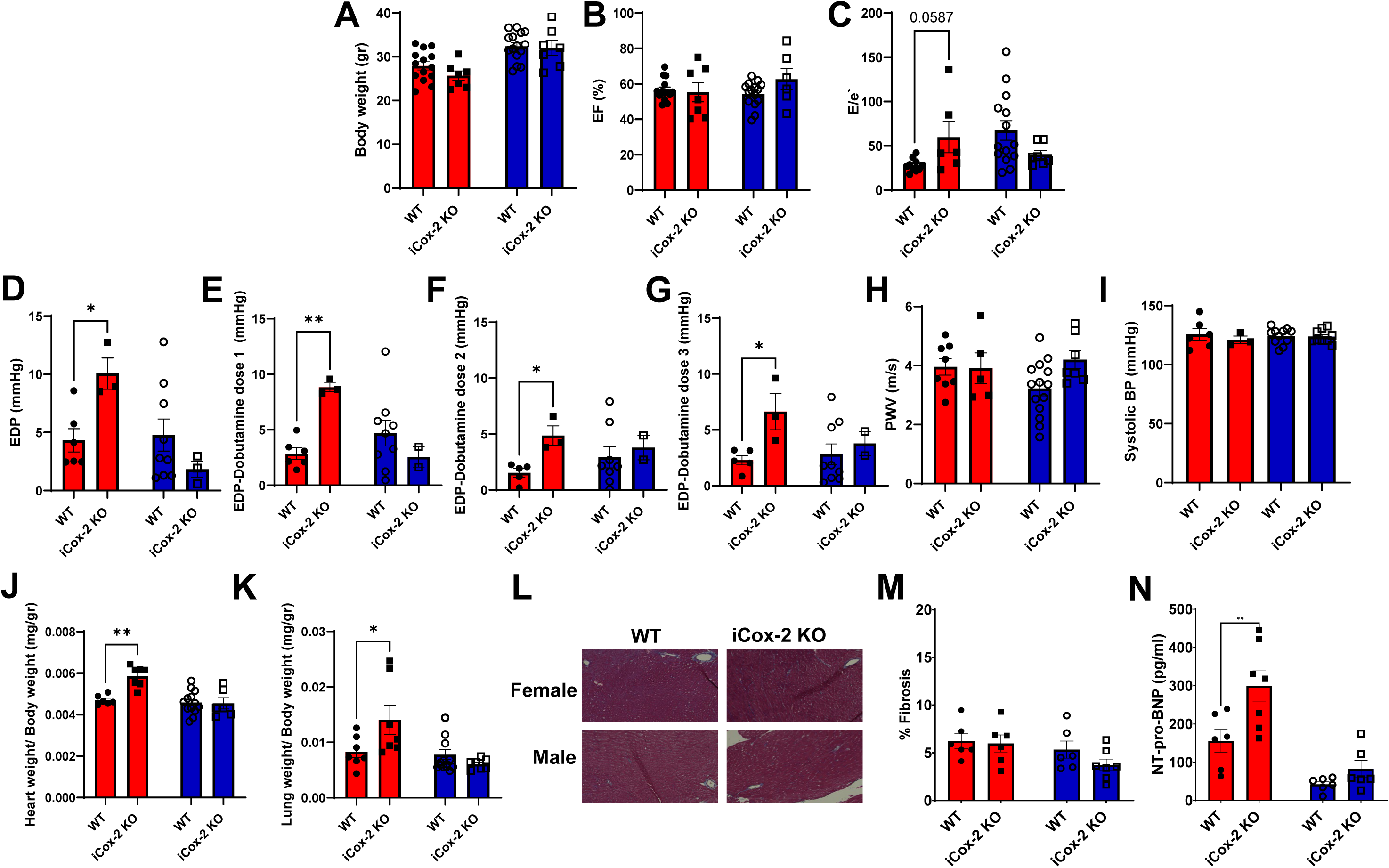
Cox-2 deletion predisposes to diastolic dysfunction in aged female mice. (A-H) Cardiac function characterization of 20-26-month-old iCox-2 KO female and male mice and sex- and age- matched WTs. A) Body weight. (B) Ejection fraction (EF). (C) Ratio of peak early Doppler transmitral flow velocity (E) to myocardial tissue Doppler velocity (e′) ratio. (D) End diastolic pressure (EDP). (E) EDP after first dose of dobutamine. (F) EDP after second dose of dobutamine. (G) EDP after third dose of dobutamine. (H) Pulse waive velocity (PWV). (I) Systolic blood pressure (BP) measured by telemetry. (J) Heart weight normalized to body weight. (K) Lung weight normalized to body weight. (L) Representative micrographs and (M) quantification of LV fibrotic remodeling because of collagen accumulation, as evaluated by picrosirius red staining. At least eight distinct non over-lapping sections were quantified to analyze the percentage fibrosis in the heart for each genotype. (N) N-terminal (NT)-pro hormone BNP (NT-pro-BNP). P values were calculated by one-way ANOVA with Sidak’s post hoc test. Bars (red= females; blue= males) and error bars show means and SEM, respectively, with individual data points superimposed.

Pulse wave velocity (PWV), a measure of vascular stiffness, was unaffected by Cox-2 deletion in both aged female and male mice (Fig. 1H, Table S6 and S8). Similarly, SBP remained unchanged in aged female and male mice following COX-2 deletion (Fig. 1I).

As compared to their sex- and age-matched WT controls, aged iCox-2 KO female mice, but not iCox-2 KO male mice, exhibited increased cardiac hypertrophy (as measured by heart/body weight ratio; Fig. 1J) and pulmonary congestion (as measured by lung/body weight ratio; Fig. 1K). Cox- 2 deletion did not affect the presence of myocardial fibrosis (Fig. 1, L-M) in male or female mice. N-terminal (NT)-pro hormone BNP (NT-pro-BNP) levels were statistically significant elevated in female but not in aged iCox-2 KO male mice (Fig. 1N).

In summary, aged iCox-2 female KO mice present features of the human HFpEF syndrome, including preserved EF (≥50%), diastolic dysfunction, cardiac hypertrophy, pulmonary congestion, and elevated NT-pro-BNP.

### Cox-2 deletion redirects arachidonic acid metabolism in aged female mice

We confirmed that aged iCox-2 female and male KO mice exhibited similar Cox-2 deletion across different organs. In aged iCox-2 female KOs, Cox-2 mRNA expression was reduced approximately 2-fold in the heart, 3-fold in the aorta, 4-fold in the renal medulla, and 2-fold in the cortex, compared to age-matched female WT mice (Fig. S2 A-D). In aged iCox-2 male KOs, Cox- 2 mRNA expression was reduced approximately 2-fold in the heart, 1.5-fold in the aorta, 4-fold in the renal medulla, and 3-fold in the cortex, compared to their age-matched male WT mice (Fig. S2 A-D). In both aged iCox-2 KO female and male mice, Cox-1 mRNA expression was unchanged by Cox-2 deletion (Fig. S2 E-H). Consistently, aged iCox-2 KOs exhibited over a 60% reduction in the urinary PGE_2_ metabolite (PGEM) and PGD_2_ metabolite (PGDM) in both female and male mice (Fig. 2A and B). The urinary metabolite of TxB_2_ (TXBM) was reduced by 50% only in aged iCox-2 KO male mice (Fig. 2C). In both female and male mice, Cox-2 deletion did not detectably reduce the already low PGI_2_ metabolite (PGIM) levels in the urine under unchallenged conditions (Fig. 2D). In aged female iCox-2 KO mice, COX-2 substrate metabolism was redirected toward 5-lipoxygenase (LOX)-dependent leukotriene (LT)E_4_ formation (Fig. 2E). Other urinary metabolites of arachidonic acid (AA), linoleic acid, eicosapentaenoic acid, and docosahexaenoic acid were not significantly altered by Cox-2 deletion in either male or female aged mice (Fig. 2F). Together, these results indicate that, while aged female and male iCox-2 KOs exhibit a similar suppression of COX-derived metabolites under basal conditions, only aged female iCox-2 KO mice show a rediversion of COX-2 substrate metabolism toward LTE_4_.

**Fig. 2.**
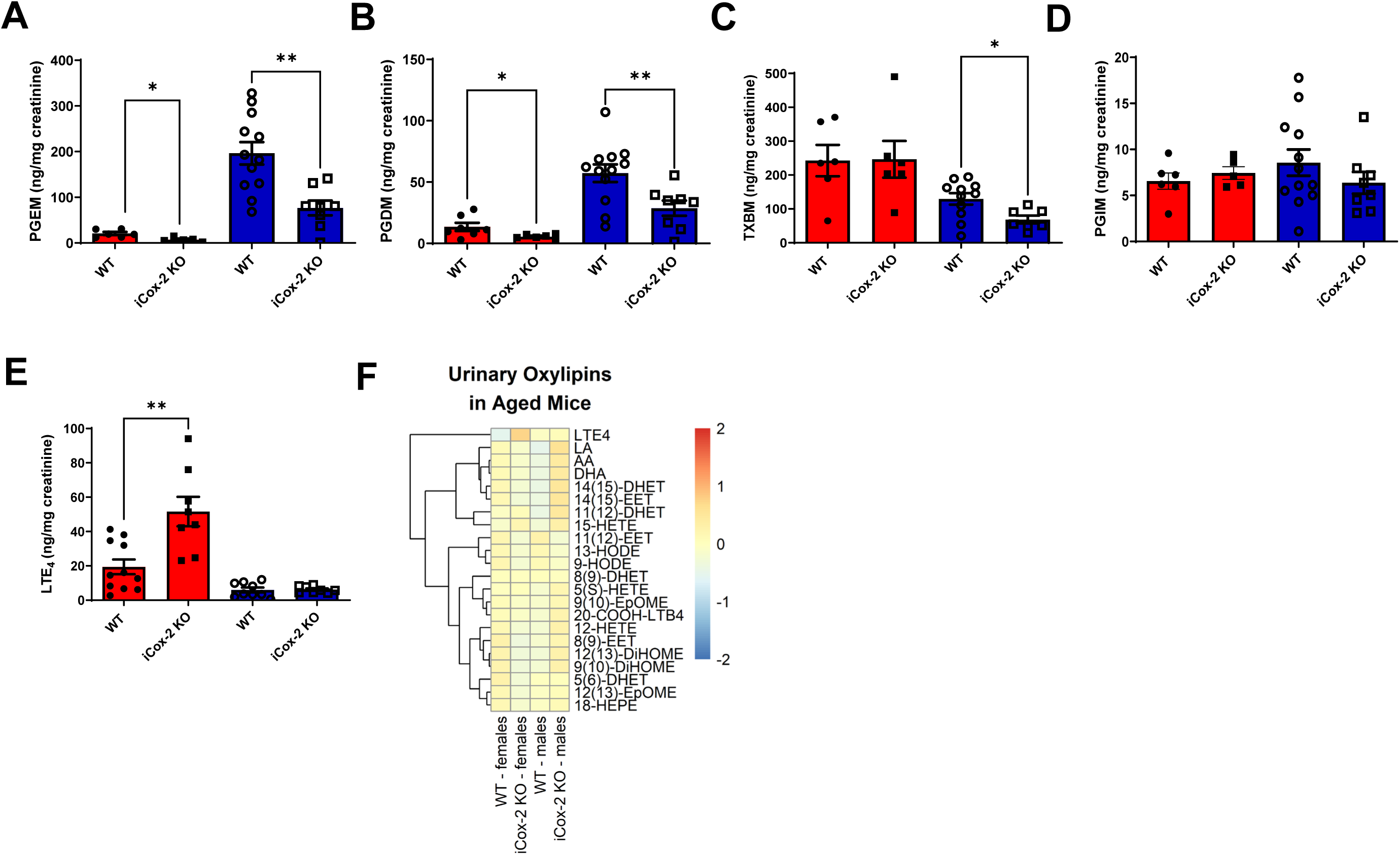
Cox-2 deletion rediverts arachidonic acid metabolism in aged female mice. Urinary metabolites of arachidonic acid, linoleic acid, eicosapentaenoic acid and docosahexaenoic acid measured in aged WT and iCox-2 KO mice by liquid-chromatography mass-spectrometry and normalized by urinary creatinine levels. (A) Urinary PGE_2_ metabolite (PGEM). (B) Urinary PGD_2_ metabolite (PGDM). (C) Urinary TXB_2_ metabolite (TXBM). (D) Urinary PGI_2_ metabolite (PGIM). (E) Urinary LTE_4_ metabolite. (F) Heatmap depicting relative levels (red = high, blue = low) of oxylipins in the urine of aged WT and iCOX-2 KO male and female mice. Pairwise *t* tests were used to determine significant differences between WT and iCox-2 KO within the same sex. Bars (red= females; blue= males) and error bars show means and SEM, respectively, with individual data points superimposed.

### Cox-2 deletion does not affect kidney function in aged mice

In the kidney, Cox-2 is expressed in both the cortex and the medulla (Fig. S2), where it regulates renal blood flow, glomerular filtration rate, and water and sodium reabsorption (29). Since kidney dysfunction occurs in approximately 50% of HFpEF patients and is associated with higher mortality (30), we examined kidney function and structure in aged iCox-2 KO female and male mice and their age-matched controls. In both aged female and male mice, Cox-2 deletion did not affect creatinine, albumin, nitrate/nitrite levels, or fractional sodium reabsorption (Fig. S3 A-F). Furthermore, the expression of genes related to the renin-angiotensin-aldosterone system (RAAS) in the renal cortex (Fig. S3 G-J) and medulla (Fig. S3 K-N) was not affected by Cox-2 deletion in aged mice. Additionally, overall kidney histology remained unaltered by Cox-2 suppression in aged mice (Fig. S3 O-R). In sum, these results indicate that, in the absence of additional stressors, Cox-2 deletion alone is not sufficient to impair kidney function in aged mice.

### Cox-2 deletion alters the expression of cardiac mitochondrial genes in aged female mice

To uncover the mechanisms by which Cox-2 deletion contributes to diastolic dysfunction in aged female hearts, we performed unbiased transcriptomic profiling of cardiac samples harvested from aged iCox-2 KO and age-matched female WT controls. RNA sequencing revealed a total of 494 transcripts differentially expressed between WT and iCox-2 KO female mice at a q-value < 0.355 (no transcripts were differentially expressed when using a lower q-value cut-off). Seventy-six percent of these transcripts were more highly expressed in the hearts of aged iCox-2 KO female mice, while twenty-four percent were more highly expressed in the hearts of aged WT female mice (Fig. 3A). Ingenuity Pathway Analysis of the differentially expressed genes between WT and iCox-2 KO mice revealed enrichment in the following pathways: oxidative phosphorylation (OXPHOS), mitochondrial dysfunction, Granzyme A signaling, Sirtuin signaling, and glutathione-mediated detoxification (Fig. 3B).

**Fig. 3.**
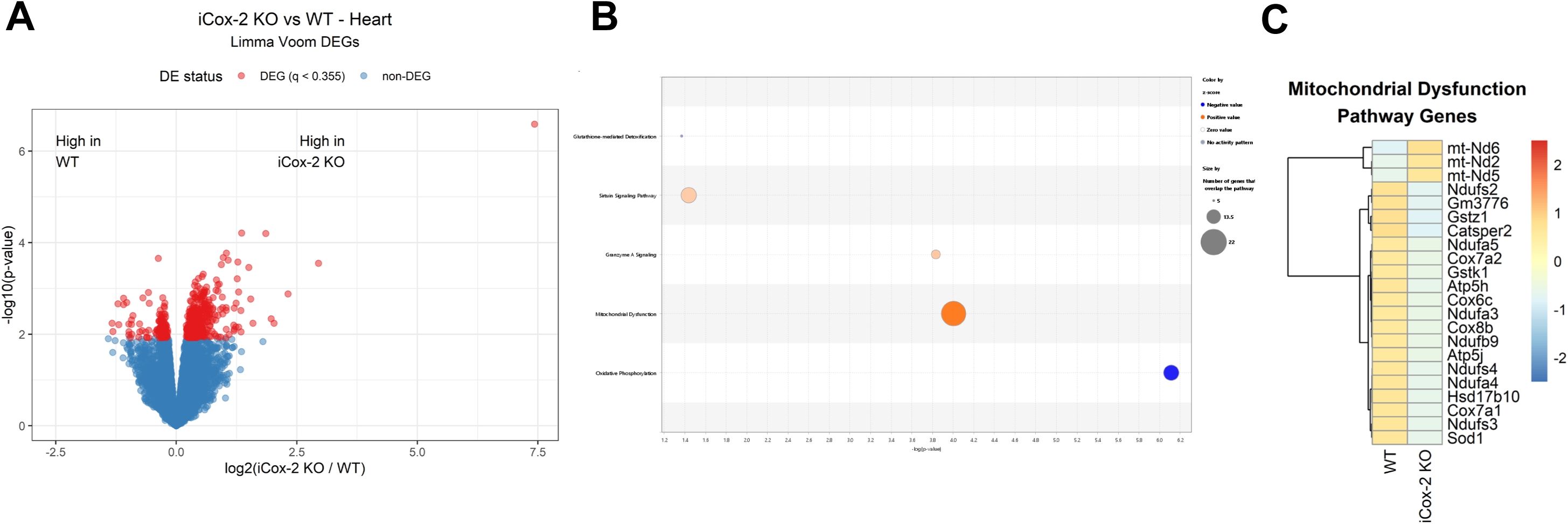
Aged iCox-2 KO female mice present cardiac muscle mitochondrial dysfunction. RNA samples isolated from the heart of aged (20-26-month-old) WT and iCOX-2 KO female mice were used for RNA-Seq (n=5 per genotype). (A) Volcano plot showing up and down-regulated transcripts in the heart of aged iCOX-2 KO female mice compared to age matched female WTs. (B) Top canonical pathways enriched by gene differentially expressed in the heart of WT and iCOX-2 KO female mice as identified by Ingenuity Pathway Analysis. (C) Heatmap depicting relative abundance of mitochondrial dysfunction transcripts differently expressed (red = high, blue = low) between aged WT and iCOX-2 KO female mice.

Mitochondrial dysfunction in cardiac muscle has been reported in patients with HFpEF (31). Among the genes in the mitochondrial dysfunction pathway, transcripts related to energy metabolism (ATP synthase, cytochrome c oxidase, NADH oxidoreductase, short-chain dehydrogenase/reductase) and antioxidant activity (glutathione transferases and superoxide dismutase) were downregulated in the hearts of aged iCox-2 KO female mice. In contrast, three transcripts encoding NADH dehydrogenase subunits were upregulated by Cox-2 deletion (Fig. 3C).

Altogether, these data are consistent with the hypothesis that mitochondrial dysfunction in cardiac muscle contributes to the HFpEF phenotype observed in aged iCOX-2 KO female mice.

### Cox-2 deletion does not impair energy metabolism or oxidative stress in aged mice

In the heart, mitochondrial dysfunction can impair energy supply, enhance oxidative stress, and disrupt calcium handling. Since disturbances in cardiac metabolism underlie most cardiovascular diseases, including HFpEF (32), we performed nontargeted metabolomics of plasma, heart, aorta, kidney cortex, and kidney medulla samples collected from aged iCox-2 KO female and male mice and their age- and sex-matched WT controls. We measured aqueous metabolites involved in various functions (amino acids, nucleotides, TCA cycle intermediates, urea cycle intermediates, and oxidative stress markers), intact lipids (including triglycerides and phospholipids), and acyl- carnitines. The heatmaps did not reveal genotype-dependent alterations in plasma and tissue metabolomic profiles in either female (Fig. S4) or male (Fig. S5) aged mice, as previously reported in adult mice (26). In summary, these results indicate that Cox-2 deletion does not affect energy metabolism or oxidative stress levels in aged mice.

### Cox-2 deletion disrupts calcium handling in the heart of aged female mice

Since we did not detect an impact of mitochondrial dysfunction on cardiac energy metabolism and oxidative stress by metabolomics, we investigated the effect of altered mitochondrial homeostasis on calcium handling in the heart. Interestingly, RNA-Seq analysis revealed that the transcript encoding phospholamban (Pln), a protein that inhibits cardiac muscle sarco(endo)plasmic reticulum Ca²⁺-ATPase (SERCA2), was more highly expressed in the hearts of aged iCox-2 KO female mice compared to age-matched WT female mice (Fig. 4A). We confirmed the differential expression of Pln between iCox-2 KOs and WTs in the hearts of aged female mice, but not in aged male mice, by RT-PCR (Fig. 4B). A similar effect was not observed in the hearts of either adult male or female mice (Fig. 4C). We then investigated the protein expression of key mediators of cardiomyocyte relaxation, specifically SERCA2 and PLN, both in their unphosphorylated and phosphorylated states. Unphosphorylated PLN inhibits SERCA2 activity and impairs cardiac relaxation by preventing calcium import into the sarcoplasmic reticulum. We found that the ratio of phosphorylated to total PLN was decreased in the hearts of aged iCox-2 KO female mice compared to age-matched WT female controls (Fig. 4 D-F). Cox-2 deletion did not change SERCA2 expression (Fig. 4G) but increased the PLN/SERCA2 ratio (Fig. 4H) in aged female hearts. Similar changes were not observed in the hearts of aged male mice (Fig. 4 D-H).

**Fig. 4.**
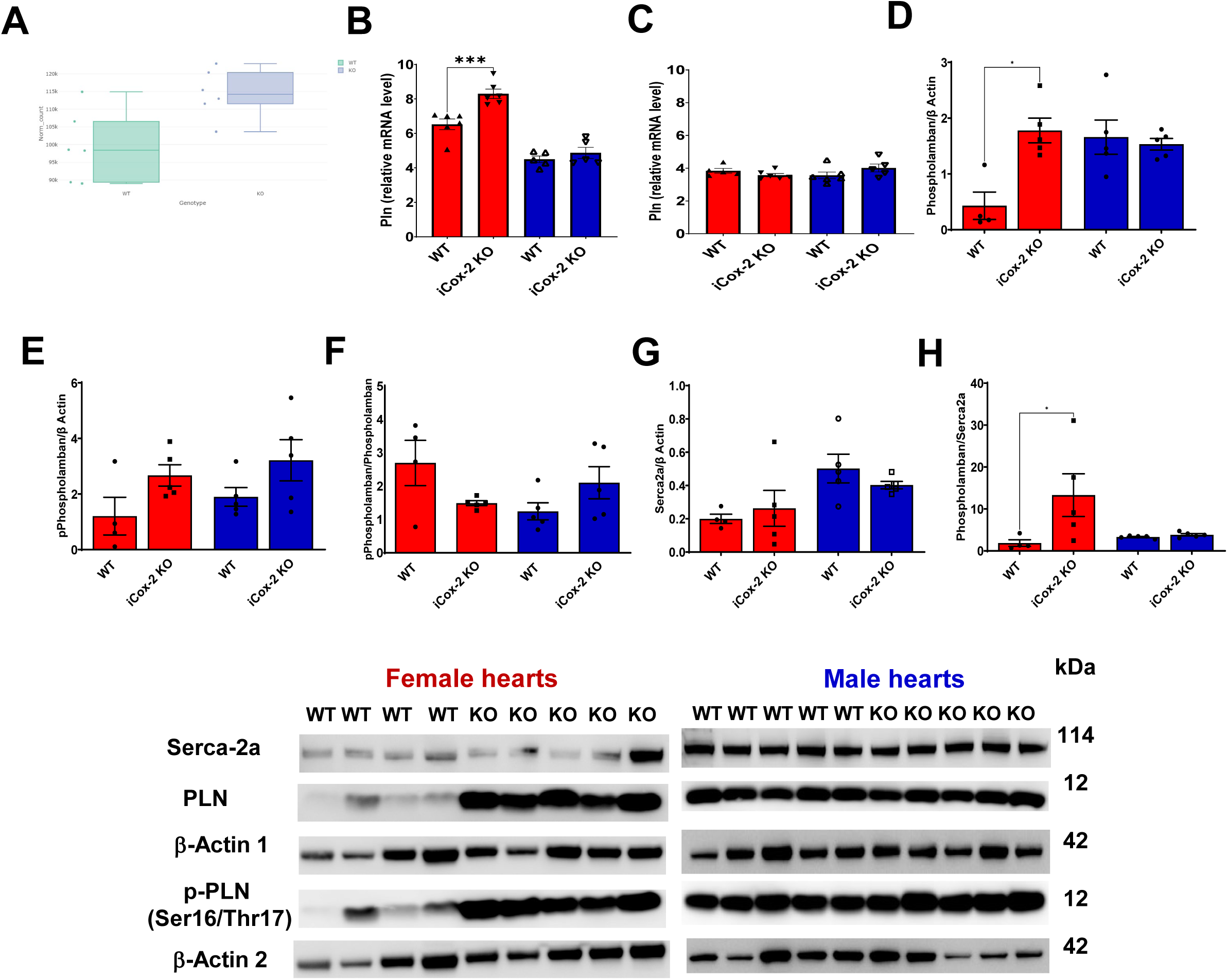
Cox-2 deletion alters calcium handling in the heart of aged female mice. (A) Number of reads mapped to Pln in RNA-Seq analysis of the heart of aged WT and iCOX-2 KO female mice. (B) Gene expression of Pln in the heart of aged WT and iCOX-2 KO female and male mice by RT-PCR. (C) Gene expression of Pln in the heart of adult WT and iCOX-2 KO female and male mice by RT-PCR. (D-H) Quantification of the protein expression of total PLN, phospho-PLN, SERCA2a in the heart of aged WT and iCOX-2 KO female and male mice measured by western blotting and quantified using the ImageJ software. Pairwise *t* tests were used to determine significant differences between WT and iCox-2 KO within the same sex. Bars (red= females; blue= males) and error bars show means and SEM, respectively, with individual data points superimposed.

In summary, these data reveal that Cox-2 deletion increases the level of unphosphorylated PLN, which can inhibit SERCA2 (increased PLN/SERCA2 ratio), and may thereby impair cardiac relaxation in the hearts of aged female mice.

### COX-2 inhibition impairs diastolic function and calcium handling in the zebrafish heart

To determine whether our findings were conserved across species, we investigated the effect of celecoxib, an NSAID selective for COX-2 inhibition, on cardiac performance in zebrafish. Both PGE_2_ and PGF_2α_, the only prostaglandins detected in zebrafish homogenates, were suppressed by celecoxib (Fig. S6), along with total cardiac output (Fig. 5A). This was not due to impaired stroke volume (Fig. 5B) but was associated with a modest, yet significant reduction in heart rate (Fig. 5C). Subsequent analysis revealed that celecoxib-treated zebrafish displayed increased relaxation time (Fig. 5D), while the time to peak contraction was marginally increased compared to vehicle treated zebrafish (Fig. 5E), indicating diastolic, but not systolic dysfunction. Fractional area change was unaltered in both celecoxib- or vehicle-treated zebrafish (Fig. 5F), however celecoxib elevated the cardiac wall stress marker, brain natriuretic peptide (*nppb)* (Fig. 5G). There were no differences in intracellular calcium transient duration during systole and diastole (Fig. 5I-N), but celecoxib increased the ventricular (Fig. 5O) and atrial (Fig. 5P) Ca^2+^ transient amplitude compared to vehicle-treated embryos. Overall, these data reveal that COX-2 inhibition in zebrafish does not affect systolic function, but impairs diastolic function and disrupts Ca^2+^ handling, as seen in aged iCox-2 female KO mice.

**Fig. 5.**
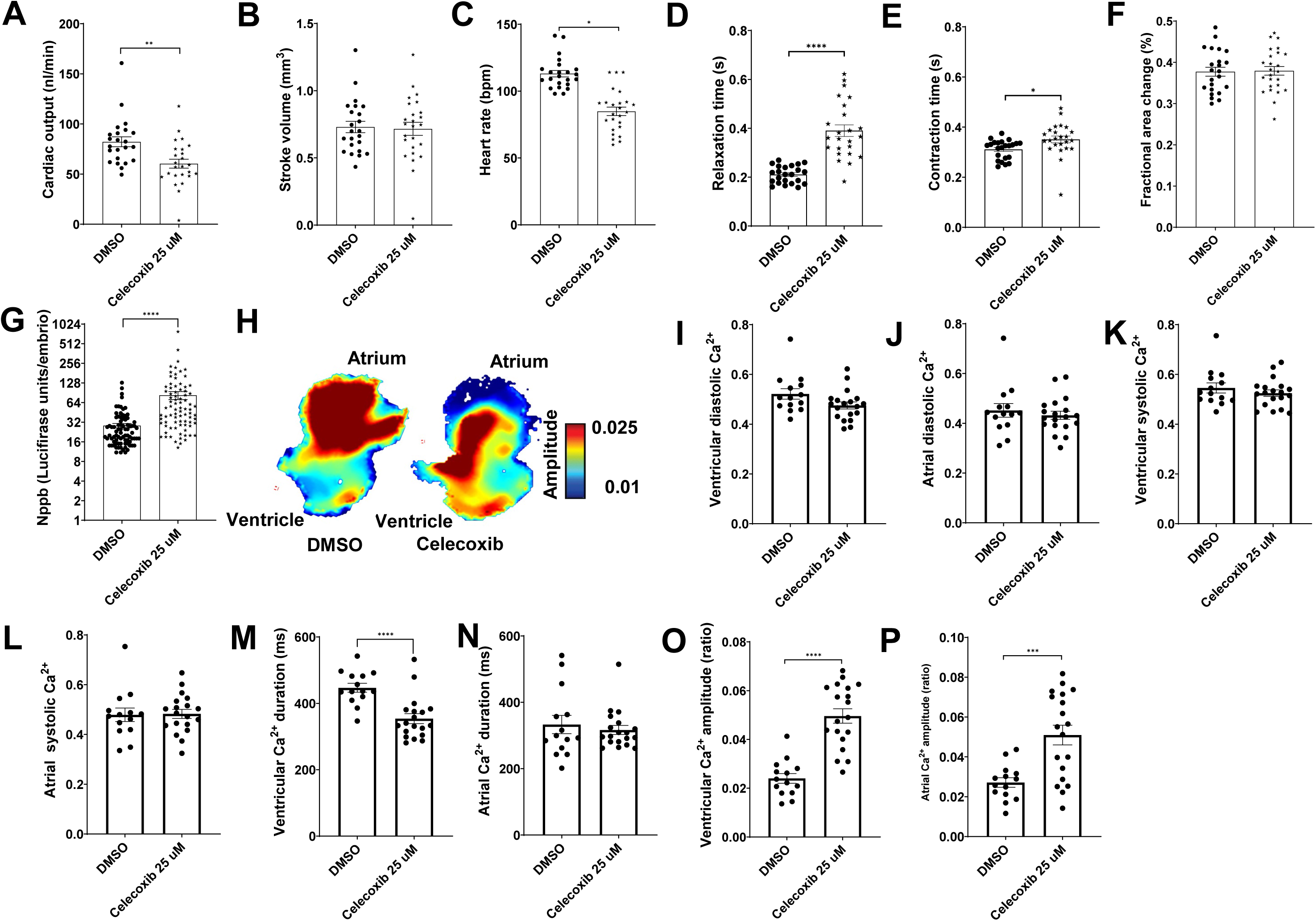
COX-2 inhibition impairs diastolic function and unbalances cardiac calcium homeostasis in larval zebrafish. Effect of celecoxib (25 μM) on cardiac performance in zebrafish larvae (A-F). (A) Cardiac output. (B) Stroke volume. (C) Heart rate. (D) Relaxation time. (E) Contraction time. (F) Fractional area change. (G) Brain natriuretic peptide (nppb) expression measured by a luciferase assay. (H) Representative color maps of calcium dynamics. (I) Ventricular diastolic calcium concentration. (J) Atrial diastolic calcium concentration. (K) Ventricular systolic calcium concentration. (L) Atrial systolic calcium concentration. (M) Ventricular calcium duration. (N) Atrial calcium duration. (O) Ventricular calcium amplitude. (P) Atrial calcium amplitude. Pairwise *t* tests were used to determine significant differences between DMSO (control) and celecoxib. Bars and error bars show means and SEM, respectively, with individual data points superimposed.

### Exposure to NSAIDs selective for COX-2 inhibition is associated with an increased risk of HFpEF in diabetic patients

To determine the clinical relevance of our findings in mice and zebrafish, we aimed to identify the type of HF associated with NSAID exposure by analyzing a subset of the Harvard-Partners electronic medical record (EMR) data, which includes 314,292 diabetic individuals. To classify the type of HF (preserved ejection fraction [EF ≥ 50%] or reduced EF [EF < 50%]), we designed an algorithm capable of extracting EF values from clinical narrative notes. In the subjects included in this study, the HF diagnosis was made after they received their first prescription for an NSAID. A manual chart review conducted by a physician confirmed that our algorithm correctly classified the HF type in 97.8% of 185 randomly selected HF patient notes.

Using a multinomial multivariable regression approach, our model revealed that, after adjusting for HF confounders, the odds ratio (OR) for HFpEF was significantly higher in both male and female subjects exposed to NSAIDs selective for COX-2 inhibition compared to those exposed to non-selective NSAIDs at 12- and 24-month follow-up (Fig. 6A). A similar trend was observed when we restricted our analysis to subjects who had not received a prescription for diuretics, commonly recommended for HF patients, before the HF diagnosis, at both follow-up periods (Fig. 6B).

**Fig. 6.**
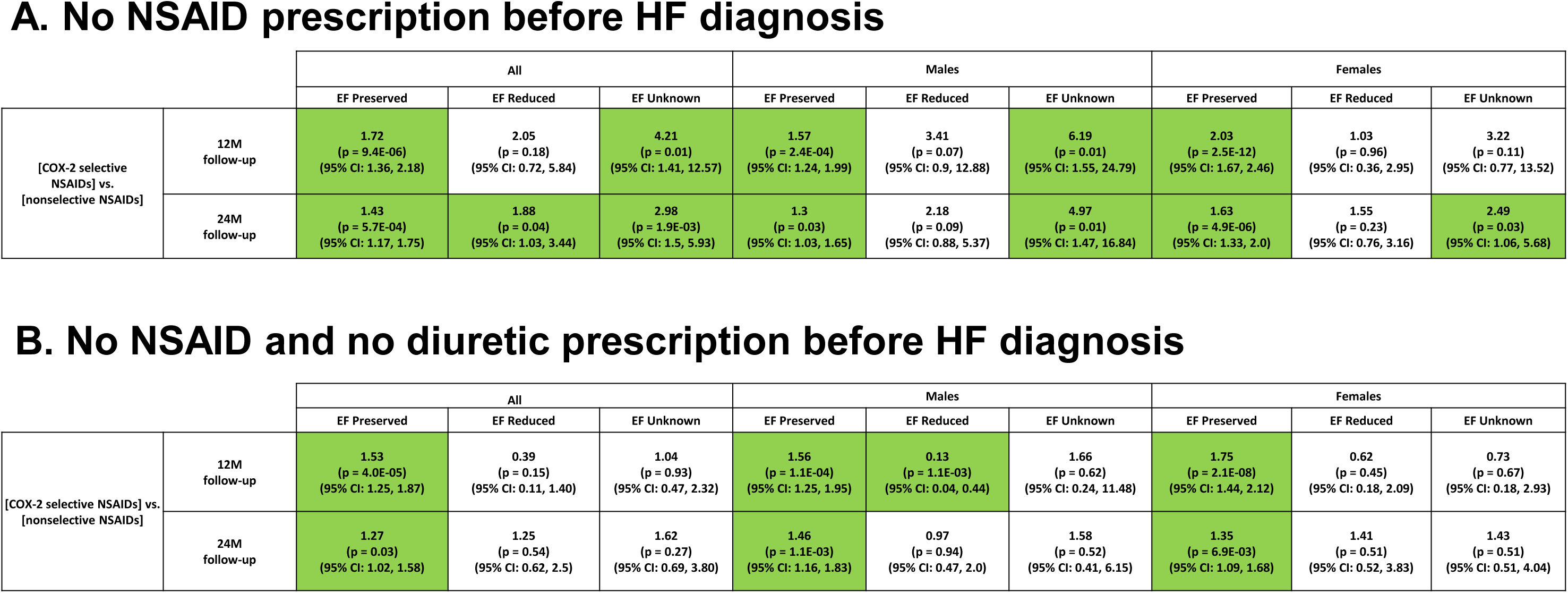
Exposure to NSAIDs selective for COX-2 inhibition is associated with an increased risk of the HFpEF in humans. (A) Adjusted hazard ratios of heart failure with preserved, reduced or unknown ejection fraction for exposure to NSAIDs selective for COX-2 inhibition vs non-selective NSAIDs in patients at 12- and 24- month follow-up after the first NSAID prescription. Ejection fraction (EF) preserved ≥ 50 %; EF reduced < 50 %. Hazard ratios (top), p-values (middle), confidence interval (bottom). Green box= p ≤ 0.05. White box = no statistical significance. (B) Adjusted hazard ratios of heart failure with preserved, reduced or unknown ejection fraction for exposure to NSAIDs selective for COX-2 inhibition vs non-selective NSAIDs in patients at 12- and 24- month follow-up after the first NSAID prescription. No diuretic prescriptions were allowed before HF diagnosis. EF preserved ≥ 50 %; EF reduced < 50 %. Hazard ratios (top), p-values (middle), confidence interval (bottom). Green box= p ≤ 0.05. White box = no statistical significance.

In summary, our model provides evidence of an association between exposure to COX-2 selective NSAIDs and an increased risk of HFpEF in humans.

## Discussion

The mechanistic link between NSAIDs and HF, as revealed by randomized clinical trials and population studies, remains remarkably under-investigated, despite representing the cardiovascular risk most strongly associated with these drugs (rate ratio 2.28, according to the CNT meta-analysis; 5). Given that NSAIDs are among the world’s most widely consumed drugs, further elucidation of this association could have significant clinical implications.

Previously, we reported that mice lacking Cox-2 in cardiomyocytes (CM-Cox-2 KO) exhibited myocardial perivascular fibrosis, abnormal contractility, exercise intolerance, arrhythmia, and preserved systolic function (34). Similarly, Cox-2 deficient rats, which share developmental defects with conventional Cox-2 KO mice (27), displayed myocardial interstitial and perivascular fibrosis, dysregulated cardiac energy metabolism, and preserved systolic function (35). Furthermore, mice treated with diclofenac, a tNSAID with selective COX-2 inhibition (13), demonstrated impaired diastolic function, but not systolic function, because of cardiac mitochondrial dysfunction (36). These preclinical studies consistently reported preserved systolic function following COX-2 deletion or inhibition, with potential diastolic dysfunction. However, the presence of extracardiac abnormalities and contributions from comorbidities were not addressed in these studies.

In this study, using a multispecies approach, we confirmed that COX-2 inhibition or deletion alone does not impair systolic function but may lead to diastolic dysfunction in both mice and zebrafish. Additionally, analysis of the Harvard-Partners clinical EMR database using a multinomial multivariable regression approach revealed an increased odds ratio for HFpEF, but not HFrEF, associated with exposure to COX-2 selective versus nonselective NSAIDs. This association was evident both when we included subjects exposed to NSAIDs before an HF diagnosis and when we restricted the analysis to those without a prescription for diuretics prior to the HF event who were exposed to NSAIDs before the HF diagnosis.

HFpEF is the most common form of HF in the elderly (37, 38), and age-related molecular and cellular alterations have been implicated in its pathogenesis (39). Female sex is associated with a higher prevalence of HFpEF (37, 40) and worse clinical outcomes (41), suggesting that sex-dependent mechanisms may contribute to its development. These mechanisms may include differences in cardiac structural adaptation to pressure overload, cardiac electrophysiology, autonomic tone, and the activation of both adaptive and innate immunity.

Our adult Cox-2 deficient mice (4–5 months old, corresponding to a human age of 40–50 years) did not exhibit any cardiac phenotype resulting from COX-2 deletion or inhibition. However, aged Cox-2 deficient female mice (20–26 months old, corresponding to a human age of approximately 70 years), but not aged male mice, displayed several hallmarks of human HFpEF syndrome, including impaired diastolic function, reduced cardiac reserve, cardiac hypertrophy, lung congestion, and elevated NT-pro BNP plasma levels.

There are discordant findings regarding fibrosis in preclinical models of HFpEF, which often require a multi-hit approach to fully replicate the human condition (42). For instance, obese ZSF1 rats, which are obese, diabetic, and hypertensive, do not exhibit myocardial fibrosis (43). Additionally, fibrosis tends to be more pronounced in patients with HFrEF than those with HFpEF (44), and the extent of fibrosis in HFpEF patients is highly variable (45), reflecting the heterogeneity of the syndrome (46). Unlike CM-Cox-2 KO mice (34), aged iCox-2 KO mice did not show increased myocardial fibrosis. It is likely that global Cox-2 deletion in adult (25) and aged mice (Fig. 1L) prevented the COX-2–dependent formation of the pro-fibrotic PGF_2α_ (47) in cardiac fibroblasts, unlike what was observed in CM-Cox-2 KO mice (34).

Inducible deletion of Cox-2 reduced PGE_2_ and PGD_2_ biosynthesis, as reflected by urinary metabolites, in both male and female aged mice. In contrast, thromboxane biosynthesis was reduced only in males compared to age- and sex-matched control mice, while PGIM remained unchanged (Fig. 2). This finding contrasts with our previous observations in iCox-2 KO mice and vascular Cox-2 KO mice. In these mice, in the context of evoked phenotypes such as atherosclerosis, thrombogenesis, or hypertension, we observed a significant reduction in PGIM that correlated with worse CV outcomes (14, 25).

Notably, only aged female iCox-2 KO mice exhibited a redirection of the AA substrate of COXs towards leukotriene biosynthesis, as reflected by urinary LTE_4_ levels. Sex differences in leukotriene biosynthesis have been observed across species, consistent with the known inhibitory effect of androgens on the assembly of the 5-LOX/5-LOX activating protein complex, which is necessary for LT biosynthesis (48, 49). Since the reduction in prostanoids due to Cox-2 deletion was observed in both aged male and female mice, while the redirection of the substrate towards LTE_4_ occurred only in aged female iCox-2 KO mice, it is likely that the decreased prostanoid biosynthesis, coupled with the concomitant increase in LTE_4_, contributes to the HFpEF phenotype. LTE_4_, along with LTC_4_ and LTD_4_, is part of the group of cysteinyl leukotrienes (cysLTs), potent pro-inflammatory mediators with bronchoconstrictor and vasoconstrictor effects, involved in the development of allergies and asthma (50). Since LTE_4_ is the most stable of the cysLTs, its levels reflect total cysLT production (51). Although the role of cysLTs in heart failure is not well established, LTE_4_ has been shown to contribute to myocardial ischemic injury by elevating Ca^2+^ levels, increasing reactive oxygen species, enhancing vascular permeability, and promoting neutrophil recruitment (52).

Mechanistically, the impact of COX-2 blockade on diastolic function appears to be mediated by myocardial mitochondrial dysfunction and impaired calcium handling. In the heart, we identified 494 differentially expressed transcripts between WT and iCox-2 KO female mice. RNA-Seq analysis revealed dysregulated pathways related to mitochondrial dysfunction in the hearts of aged iCox-2 KO female mice compared to age-matched WT female controls. The alteration in the mitochondrial transcriptional profile did not affect energy metabolism or oxidative stress at rest, but it did impact intracellular Ca^2+^ dynamics. PLN was upregulated in the hearts of aged female iCox-2 KO mice at both the gene and protein levels. Human genetics studies of dilated cardiomyopathy and genome-wide association studies have shown that variation in PLN is associated with HF (53–58). Cox-2 deletion did not affect the expression of SERCA2a or the phosphorylation rate of PLN. In aged female iCox-2 KO mice, the PLN/SERCA2a ratio was increased, indicating a stronger interaction between PLN and SERCA2a, reduced SERCA2a affinity for Ca^2+^, and, consequently, impaired myocardial relaxation.

Zebrafish treated with celecoxib, at a dose that impairs diastolic but not systolic function and increases Nppb expression, also exhibited Ca^2+^ mishandling, characterized by an increase in the amplitude of calcium transients. Similarly, HFpEF rats showed augmented calcium transient amplitudes and increased PLN expression in their cardiomyocytes (59).

In this study, the diastolic dysfunction observed in aged iCox-2 KO female mice was not due to alterations in hemodynamics or renal function resulting from Cox-2 deletion in the kidney. This is despite kidney dysfunction being a non-cardiac comorbidity associated with HFpEF (30) and NSAIDs having the potential to alter water and sodium retention, thus predisposing to hypertension (29). In the absence of additional stressors, Cox-2 deletion alone was insufficient to impair renal function in aged mice, and iCox-2 KO mice were not hypertensive. This is consistent with our previous work, in which we reported that only celecoxib-treated mice infused with angiotensin II at a dose sufficient to elevate BP showed impaired renal function and increased formation of methylarginines in the kidney (26).

In summary, our EMR study revealed that NSAIDs selective for COX-2 inhibition increase the risk of developing HFpEF compared to HFrEF. Similarly, our preclinical studies suggest that COX-2 deletion or inhibition does not impair systolic function but may affect diastolic function. Future studies should investigate the effects of COX-2 deletion or inhibition on cardiac function in the presence of metabolic disease, obesity, and/or hypertension better to reflect the comorbidities common in regular NSAID users. These studies should also further explore the contributions of age and sex.

In conclusion, we provide initial evidence suggesting that perturbation of the COX-2 pathway may be one of the multiple mechanisms involved in the HFpEF syndrome. Impaired calcium handling appears to contribute to the myocardial relaxation deficits associated with COX-2 disruption. Restoring Ca²⁺ homeostasis could therefore offer a promising strategy to mitigate the risk of heart failure in patients taking NSAIDs.

### Methods summary

The aim of the EMR study was to investigate the type of HF, classified as EF ≥ 50% or EF < 50%, associated with COX-2 selective or nonselective NSAID exposure by analyzing large human observational databases using a multinomial multivariable regression approach.

The aim of the studies in model organisms was to investigate the effect of COX-2 deletion and inhibition on cardiac function. In mice, we characterized cardiac function using echocardiography, invasive hemodynamics, and blood pressure measurements via telemetry. Mechanistic studies included RNA-Seq, urine lipidomic profiling, and metabolome profiling of plasma, heart, aorta, and kidney, as well as Western blotting. In zebrafish, we assessed cardiac performance, measured Ca^2+^ dynamics, and evaluated nppb expression using a luciferase assay. Inhibition of prostaglandin biosynthesis *in vivo* and plasma celecoxib levels were measured by mass spectrometry.

Sample sizes were empirically determined based on our previous experience with the animal models used (14, 25–26, 34). All animals were randomly allocated to treatment groups, and the experimenters were blinded to the allocation. The exact number of animals, along with their sex and age, for each experiment is provided in the respective figures. Please refer to the Supplementary Materials and Methods for further details on experimental procedures.

### Statistical analysis

Data are presented as bar with error bars showing means and SEM, respectively, with single data points superimposed. Indicated sample sizes in figure legends refer to biological replicates. Comparisons between two groups were done by Mann-Whitney’s U test. In the case of multiple group comparisons, analysis of variance (ANOVA) followed by Sidak’s post hoc tests were applied.

Statistical analysis of EMR data and omics data are described in detail within their respective methods sections in Supplementary Materials and Methods. All reported P values are two-sided, and an α level of 0.05 was used throughout.

Unless stated otherwise, statistical analyses were performed using R package (version 3.6.0) or GraphPad Prism 10 (GraphPad Software LLC, Massachusetts, USA).

## Funding

This study was supported by a grant from the NIH (HL117798).

## Author contributions

ER and PH designed and supervised the study; ER, and GAF wrote the manuscript; ER, MB, CC, SYT, SG, USD, JA, NJL, TW, performed experiments and analyzed and discussed data; UK and SYS interrogated and analyzed the clinical data in EMRs; ER, PH, JLG, SYS, CAMR, GAF discussed data and/or gave conceptual advice.

## Competing interests

G.A.F. is the McNeil Professor in Translational Medicine and Therapeutics and Senior Advisor to Calico Laboratories. All authors declare no competing non-financial interests. Data and materials availability: All data associated with this study are present in the paper or the Supplementary Materials. The RNA-Seq data are accessible on the NCBI GEO database under the accession number GSE281720.

## Materials and Methods *Human Study*

We used Partners HealthCare’s EMR database, a repository that contains data entries collected at two large tertiary care academic hospitals, Massachusetts General Hospital (MGH) and Brigham and Women’s Hospital (BWH), according to institutional review board–approved protocols. Of the entire patient population of over 4 million patients, we focused on those who received care between 1990 and 2010 and who tend to highly utilize medical resources. For this reason, we selected a diabetic sub-population with individuals who had any of the following criteria: ≥1 ICD9 for diabetes, ≥1 diabetic medication, HbA1c ≥ 6.5%, or plasma Glucose ≥ 200 mg/dl.

These selection criteria yielded a set of 314,292 patients. We applied to this population an innovative case-control study design. As cases (“exposed”), we identified patients who were prescribed at least two NSAIDs selective for COX-2 inhibition separated by at least 30 days, reassuring a continuous use of the drugs (Table S9). As controls (“unexposed”), we identified patients that were prescribed two or more of nonselective NSAIDs throughout their lifetime, similarly, to reassure a continuous use of the drugs (Table S10).

We defined the baseline time step as the date in which an exposed patient was prescribed: a) COX-2 selective NSAID for the first time throughout his or her lifetime; b) an unexposed matched patient was an individual who was at a similar age as of the exposed patient at baseline, as defined by the following bins, given in years: [18–25], [25–30], [30–35], … , [80–85], [85– 90], and [90+]. Additionally, each one or more unexposed patients matched to an exposed patient had a similar profile of utilization of care as described further in this section. An additional criterion to define unexposed patients was to restrict them from having any COX-2 selective NSAID prescriptions prior to baseline of their matched exposed patients. To match exposed and unexposed patients based on care utilization, we defined the notion of “EMR facts”; a fact was defined as any data entry found in the Partners system that was associated with the patient including labs, medications, any type of encounter, etc. Unexposed were matched with exposed patients for number of EMR facts over the patient’s lifetime. The larger the number of facts, the higher the utilization of care consumed by the patient as defined by the following bins: [100– 500], [500–1,000], [1,000–2,000], [2,000–3,000], [3,000–4,000], [4,000–5,000], [5,000+].

For all exposed and unexposed patients, we extracted demographic confounders (age, gender and race) in addition to multiple comorbidities known to impact HF or medication status (hypertension, myocardial infarction, coronary heart disease, renal failure, joint disorders, and GI disorders, Table S11). Furthermore, for all exposed and unexposed patients we observed 12- and 24-months follow-up periods after baseline at two types of possible outcomes including: HFpEF (EF≥ 50%) or HFrEF (EF< 50%).

To assure continuous care prior to baseline we filtered out all patients with no interaction with the care system at least 12 months before the baseline (e.g., such patients had no indication for details such as office visits, vital signs, laboratory observations). We made sure that all patients had neither inpatient nor outpatient HF ICD-9-CM codes in their EMR profile prior to baseline. To identify if a patient was associated with HF during the follow-up period, we observed if the patient had either an inpatient ICD-9-CM code of 428.xx or an inpatient diagnosis-related-group (DRG) DRG:127 code (*60*). Additionally, to reassure reliability of billing codes for HF, we considered a billing code as a true indication for HF only if it was associated with a diuretic code prescription (Table S.12) ±7 days within the date of the HF. Given that a certain patient had an inpatient HF code during the follow-up period and a prescription of diuretic, we determined the type of the HF by observing left ventricular EF one or more measurements that were associated with each patient (both exposed and unexposed patients). In Partners’ EMR, EF values are stored in free-text reports such as echocardiographic, radiology, and discharge reports. To extract the EF values from such unstructured resources we developed an extraction algorithm that handled the heterogeneous documentation styles of physicians at MGH and BWH. For example, we extracted EF = 45% for “The estimated left ventricular ejection fraction is 45-50%”; EF = 67% for “An echocardiogram on 6/7 showed an ejection fraction of 67 percent” and EF = 42% for “… LVEF 42%”. In total the algorithm extracted from the various reports approximately 0.5-million EF values representing our initial 314,292-patient population. We decided to consider echocardiogram and radiology reports for extraction of EF values, but not discharge summaries as discharge summaries contain often multiple EF values, a mixture of historical and current values, which makes the dates associated with the extracted EF values difficult to assess.

If a patient had multiple inpatient HF codes in different dates, then we considered the first occurrence as the date of the HF. To identify the type of HF, we considered then the EF measurement found at the same day of the HF code or the earliest subsequent after the HF code and within the follow-up period. We ignored EF measurements that occurred before the HF or any other EF measurements. In rare cases in which we were not able to find an EF value in the period between the date of the HF code and the end of the follow-up period, we considered the EF value between the baseline and the HF code date. Additionally, we filtered out patients who had no HF within the follow-up period and had no recorded interaction with the care system after the follow-up period. The logic behind that was to make sure that the patients that we considered received continuous care either in MGH or BWH and to be sure that they truly had no HF during the follow-up period and did not die or were alive but admitted for HF in a different hospital.

Our study did not have the advantage of random treatment assignment, and as such we implemented across our entire experiments a multinomial multivariable regression incorporating a propensity score matching methodology (*61*). Specifically, we followed the inverse probability treatment weighting (IPTW) approach in which the contributions of the study subjects are weighted. The weights assure that for each combination of baseline characteristics, the sum of contributions of all experimental and control patients are equal. Thus, IPTW generates a pseudo-population in which each covariate combination is perfectly balanced between treatment groups (*62*). To calculate standard errors and *P* values we used bootstrapping with 1,000 re-samples.

### Mouse study

C57BL/6J mice were purchased from the Jackson Laboratory. Tamoxifen-inducible Cox-2 KO (iCox-2 KO, COX-2^flox/flox^ CreER^T2^ ^+/-^) mice and their littermate controls (WT; COX-2^flox/flox^, Cre CreER^T2-/-^) were generated as previously reported (*25*). Tamoxifen (100 mg/Kg) was injected i.p. for 5 consecutive days in iCox-2 KO mice and their controls at 8 weeks of age. All the experiments were performed at least 4 weeks after tamoxifen injection. ICox-2 knockout mice and their controls were fully backcrossed on C57BL/6.

In all cases, C57BL/6J mice and transgenic deficient mice in the indicated gene were compared with appropriate strain-, age-, and sex-matched control animals. All the experiments were performed in adult mice (4-5 month old) and in aged mice (20-26 month old).

C57BL/6J mice, 2-3 month old, received a diet containing celecoxib (100 mg/kg body weight per day in PMI rodent diet #5001), rofecoxib (50 mg/kg body weight per day in PMI rodent diet #5001), or a control diet (PMI rodent diet #5001) for 8 weeks as previously reported (*26*). Celecoxib and rofecoxib were obtained from Sequoia Research Products. Drug-containing diet was prepared by Teklad laboratory.

From these mice, blood was collected from the vena cava and, after centrifugation, plasma was separated for lipidomic analysis, metabolomic analysis, for the assessment of the kidney function or drug plasma concentration. Heart, aorta, kidney cortex and medulla were harvested and used for real-time polymerase chain reaction analysis of gene expression.

The investigator assessing phenotype was unaware of the genotype throughout these experiments. All animals in this study were housed following guidelines of the Institutional Animal Care and Use Committee (IACUC) of the University of Pennsylvania. All experimental protocols were IACUC-approved.

#### 2D Tissue Doppler Echocardiography

A comprehensive transthoracic 2-dimensional (2-D) and Doppler echocardiographic study was conducted using a high-resolution (30-MHz) Vevo 770 imaging system (VisualSonics Inc., Toronto, Canada) as previously described (*63*). The parasternal long- and short-axis views were used to record 2D images and to guide M-mode recordings obtained at the mid-ventricular level M-mode images and real time 2D B mode cine-loops of short- and long-axis views of the left ventricle were recorded and analyzed for cardiac structure and function assessment. The transducer was placed along the long-axis of the LV and directed to the right side of the neck of the mouse to obtain 2D LV long-axis views. The transducer was then rotated clockwise by 90 degrees and the LV short-axis was obtained. 2D guided LV M-mode at the papillary muscle level was recorded in the short-axis view. Cardiac structural remodeling and systolic function parameters were collected including wall thickness and LV diastolic and systolic dimensions. Pulsed Doppler studies of LV diastolic function were performed in the apical 4-chamber view with the Doppler cursor oriented parallel to the long-axis plane of the LV. Trans-mitral inflow color Doppler spectra was recorded by placing the sample volume just below the level of the mitral valve annulus and adjusted to capture the highest early diastolic flow velocity peak of the transmitral Doppler flow signal. The early and late diastolic peak velocity (E, A) and their ratio (E/A) were derived from the transmitral Doppler waveform. LV systolic intervals of the isovolumic contraction time (IVCT), the LV ejection time, IVRT, the acceleration and deceleration times of the E-wave (EAT and EDT) were derived from the waveform.

#### Invasive Hemodynamics

Invasive hemodynamic recordings were conducted as previously described (*64*). Briefly, mice were anesthetized using 2% isoflurane. Buprenex was used to alleviate pain during surgery. Mice were placed on a heating pad and a rectal temperature probe was used to maintain body temperature between 36° C and 37° C. A surgical approach was employed to expose the right carotid artery. Proximal aortic and LV intracardiac pressures were recorded with a 1.4F microtip catheter (SPR-839; Millar Instruments, Houston, TX). The catheter was zeroed in a saline bath before acquiring measurements. The catheter tip was inserted into the artery and advanced retrograde down the ascending aorta through the aortic valve into the LV. Signals were digitized at 2kHz using PowerLab/16 SP A/D converter (AD Instruments Ltd, Mountain View, CA) and stored on a PC hard drive. Analysis was performed in LabChart. Following placement of the catheter in the LV, the mice were allowed to stabilize for 5-10 minutes and to warm up to body temperature (37°C). Measurements of intra-aortic pressures (systolic, diastolic, and mean) were made. LV pressure waveforms were measured and used to calculate readouts for systolic function (maxdp/dt) and markers of diastolic function including mindp/dt, Tau, left ventricular end-diastolic pressure.

#### Blood pressure measurement by radiotelemetry

The implantation of radiotelemetry (model No. TA11PA-C20, Data Sciences International, St. Paul, MN) was performed as previously described (*14*). After 1 week of recovery after surgery, blood pressure was monitored continuously for three days using the Dataquest LabPRO Acquisition System.

#### Real-Time Polymerase Chain Reaction Analysis of Gene Expression

Total RNA from heart, aorta and kidney samples was isolated using the Qiagen RNeasy Kit. Reverse transcription was performed using an RNA-cDNA kit (Applied Biosystems, Carlsbad, CA). Real-time polymerase chain reaction was performed using ABI Taqman primers and reagents on an ABI Prism 7500 thermocycler according to the manufacturer’s instructions. The following primers were used: Cox-2 (Mm00478374_m1), Cox-1 (Mm00477214_m1), Ace2 (Mm01159003_m1), Agtr1a (Mm01166161_m1), Agtr1b (Mm01701115_m1), Agtr2 (Mm01545399_m1), Pln (Mm04206542_m1). All mRNA measurements were normalized to hypoxanthine guanine phosphoribosyl transferase-1 (Hprt1, Mm01545399_m1) mRNA levels.

#### RNA-Seq Analysis of Gene Expression

100 ng of total RNA from heart samples were prepared for sequencing using the Illumina TruSeq Stranded mRNA preparation kit (Illumina, Sand Diego, CA). The resulting libraries were sequenced in a paired-end 100bp read configuration on an Illumina NovaSeq. STAR v2.7.6a (*65*) was used to align the raw reads against the GRCm38 build of the mouse reference genome. The STAR alignment was run with the following command-line parameters: “--outSAMtype BAM Unsorted --outSAMunmapped Within KeepPairs --outFilterMismatchNmax 33 -- seedSearchStartLmax 33 --alignSJoverhangMin 8”. These data were normalized and quantified at the gene-level using v0.8.5e-beta of the Pipeline Of RNA-Seq Transformations (PORT; https://github.com/itmat/Normalization). Both STAR and PORT were provided with transcript models from v102 of the Enseml annotation (*66*). Differential expression analysis was performed in v4.4.0 of R using v3.60.0 of the limma package (*67*). Heatmaps and other visualizations were performed in R using v1.0.12 of the pheatmap package and v3.5.1 of the ggplot2 package, respectively.

#### Assessment of kidney function

Albumin, urea, nitrate/nitrite and sodium levels were measured using the following kits: albumin elisa kit (Abcam#108792); nitrate/nitrite colorimetric assay (Cayman# 780001); urea assay kit (Abcam #83362); sodium assay kit (Abcam #211096).

#### Fibrosis staining of the heart

Mouse samples were fixed overnight in 4% paraformaldehyde and dehydrated through an ethanol series. All samples were paraffin-embedded and sectioned at a thickness of 5 μm. Masson’s trichrome staining was completed using a standard protocol (Abcam). Slides were digitally scanned at 20× (Keyence, BZ-X810) and analyzed via color deconvolution using ImageJ software. Quantification of staining was completed by determining the ratio of positively stained (red) pixels to the total pixel number of each section (% fibrosis) by a blinded observer. Six distinct non over-lapping sections were quantified to analyze the percentage fibrosis in the heart for each genotype. Heart sections from each group were processed in parallel, and images were acquired with identical acquisition parameters. The analysis was carried out using Image-Pro Plus. Collagen content was expressed as fraction (percentage) of the total area.

#### Hematoxylin and eosin staining of the kidney

The hematoxylin-eosin (H&E) stain was performed on 5 μm sections of the formaldehyde-fixed and paraffin-embedded kidney samples according to the manufacturer’s protocol (Abcam).

The H&E images were taken using light microscopy (Keyence BZ-X810 microscope) supplied with a digital camera under a magnification of 4 X.

#### N-Terminal Pro-brain natriuretic peptide (NT-ProBNP) measurement

NT-ProBNP was measured by Elisa in heparinized mouse plasma following manufacturer’s instructions (Abbexa).

#### Mass Spectrometric Analysis of Prostanoids in urine samples

Urinary prostanoid metabolites were measured by liquid chromatography/mass spectrometry as previously described (*14*). Mouse urine samples were collected by using metabolic cages over an 8-hour period. Results were normalized with urinary creatinine.

#### Mass Spectrometric Analysis of oxylipins in urine samples

Urinary metabolites were measured by ultra high-pressure liquid chromatography/tandem mass spectrometry (LC/MS) using stable isotope-labelled internal standards as previously described (*68*). The levels of arachidonic acid (AA), linoleic acid (LA), docosahexaenoic acid (DHA), eicosapentaenoic acid (EPA), 14(15)-DHET, 11(12)-DHET, 8(9)-DHET, 5(6)-DHET, 14(15)- EET, 11(12)-EET, 8(9)-EET, 15-HETE, 12-HETE, 5-HETE, LTB_4_, LTE_4_, 20-OH-LTB4, 13- HODE, 9-HODE, 18-HEPE, 12(13)-DiHOME, 9(10)-DiHOME 12(13)- Epome and 9(10)-Epome were measured in each sample. Results were normalized with urinary creatinine.

#### Mass spectrometric analysis of plasma or urinary creatinine

Quantitation of plasma or urinary creatinine was performed using ultra high-pressure liquid chromatography/tandem mass spectrometry with positive electrospray ionization and multiple reaction monitoring as previously described (*14*).

#### Mass spectrometric analysis of celecoxib and rofecoxib plasma level

Plasma samples were spiked with internal standards and purified by solid-phase extraction (*26*). The analysis was performed on a Waters ACQUITY UPLC system in-line with a Waters Xevo TQ-S Triple Quadrupole Mass Spectrometer. The transition for celecoxib was *m/z* 380.0>316.2 and for d7-celecoxib was *m/z* 387.0>323.2. The transition for rofecoxib was *m/z* 315>269 and for d5-rofecoxib were *m/z* 320>273. All were monitored in negative mode. Quantitation was done by peak area ratio, and results were normalized to the sample volume.

#### Metabolomic analysis of the mouse heart, aorta, kidney and plasma samples

Metabolites were extracted using the methanol/chloroform/water method described previously (*26*). Briefly, 100 μL of plasma or 50 µg of tissues was added to 600 μL methanol/chloroform/water (2:2:1; vol/vol), and the samples were homogenized with a Tissue lyser (Qiagen, UK) and sonicated. Water and chloroform were added to the samples before being centrifuged. The resulting aqueous and organic phases were separated from the protein pellets. The extraction procedure was repeated on the remaining protein pellets. Both organic and aqueous phases were collected and evaporated to dryness. The dried samples were stored at −80°C until further analysis.

Half of the aqueous extract was reconstituted in 200 μL of 10 mmol/L ammonium acetate containing universally ^13^C- and ^15^N-labeled glutamic acid (Cambridge Isotope Laboratories) at a concentration of 20 μmol/L. Chromatographic separations were performed using a C18 pentafluorophenyl UPLC column (Ace) kept at 30°C on an Ultimate 3000 UHPLC (Dionex), coupled with a TSQ Quantiva (Thermo Fisher Scientific) triple quadrupole mass spectrometer.

The temperature of the autosampler was set to 7°C. The mobile phase consisted of solvent A: 0.1% formic acid in HPLC grade water (Sigma-Aldrich) and solvent B: 0.1% formic acid in acetonitrile (Sigma-Aldrich). When eluting the column, the following gradient was used: initial conditions were 100% A held for 1.5 minutes followed by a linear gradient with increase of B to 100% at 4.5 minutes with re-equilibration for 1.5 minutes, giving a total run time of 6 minutes with a flow rate of 400 μL/min. The injection volume was 2 μL. Samples were analyzed using a multiple reaction monitoring approach. The ionization mode used by the mass spectrometer was electrospray ionization with a capillary voltage of 3.5 kV for positive ion mode and 2.5 kV for negative ion mode. All compound dependent parameters were established using the Quantiva automatic optimization protocol infusion with standards of the relevant compounds.

For the acyl-carnitines analysis, half of the organic and half of the aqueous plasma extracts were recombined and dried down and 100 μL of deuterated internal standard (C0D9, C2D3, C3D3, C4D3, C5D9, C8D3, C14D9 and C16D3) in acetonitrile:water 1:1 was added. Samples were vortex mixed, sonicated (10 min) and the supernatant was transferred to 1.2 μm 96-well Filter Plates (Corning) over PlateOne 96-well Microplates with V bottoms (Starlab) for collection. The samples were centrifuged (10 min at 2000 rcf) to filter, the filter plate discarded and the collection plate sealed.

Chromatographic separations were performed using a T3 ultra-performance liquid chromatography (UPLC) column (Waters) kept at 30⁰C on an Ultimate 3000 ultra-high performance liquid chromatography system (UHPLC) (Dionex), coupled with a TSQ Quantiva (Thermo Scientific) triple quadrupole mass spectrometer. The temperature of the autosampler was set to 7⁰C. The mobile phase consisted of solvent A: 0.1 % formic acid in water and solvent B: 0.1 % formic acid in acetonitrile. When eluting the column the following gradient was used: Initial conditions were 95 % A held for 1 min followed by a linear gradient with increase of B to 100 % at 6 min, held for 2 min, with re-equilibration for 2 min giving a total run time of 10 min with a flow rate of 500 μL/min. The needle wash consisted of 10 % aqueous acetonitrile and was set at 200 μL, whilst the injection volume was 2 μL. Samples were analysed by detecting the precursors of product ions with an m/z of 85 using a multiple reaction monitoring (MRM) approach. The ionisation mode used by the mass spectrometer was electro spray ionisation with a capillary voltage of 3.5kV for positive ion mode (*69*).

For quantitative analysis, acquired data were processed with Xcalibur 2.0 (Thermo Fisher Scientific). Peaks of each analyte and internal standard were detected and integrated. Each analyte was normalized to the internal standard. 56 analytes were measured in plasma and 66 analytes were measured in the heart, aorta, and kidney samples.

### Zebrafish study

#### Cardiac performance of zebrafish hearts

Embryos were laterally positioned in a depression slide and allowed to acclimate to room temperature prior to imaging. Video microscopy was performed with a Zeiss Axioplan upright microscope, equipped with a FastCam-PCI high speed camera (Photron). A total of 1,088 frames were captured and processed further in ImageJ and Excel to analyze stroke volume, heart rate and cardiac output.

#### Luciferase Assay

3 Day-old embryos *Tg(nppb:luc)* treated with vehicle or celecoxib were placed in a 96-well plate in a volume of 100 μL and luciferase activity was measured after addition of firefly luciferase reagent as previously described (*70*) on a luminescence plate reader (Victor X, Perkin Elmer).

#### Measuring Ca^2+^ dynamics in embryonic zebrafish hearts

Hearts were isolated from 72-hour post fertilization zebrafish embryos in HEPES-buffered Tyrode’s solution and loaded with 50 mM of the ratiometric, calcium-sensitive dye Fura-2 AM (Life Technologies) for 20 min. Hearts were then transferred to and incubated in fresh Tyrode’s buffer for 45 minutes to allow complete hydrolysis of the esterified dye. Hearts were placed in an incubation chamber containing Tyrode’s solution supplemented with cytochalasin D to prevent cardiac contraction, mounted on an inverted microscope (TE-2000, Nikon). Ratiometric (380 nm/340 nm) calcium recordings of paced hearts were performed as previously described (*71*) and collected images were further processed in MATLAB using in-house developed scripts.

#### Mass Spectrometric Analysis of prostanoids and celecoxib in zebrafish

Prostanoid and celecoxib were measured using high-performance liquid chromatography–tandem mass spectrometry. Methanol extracts from homogenized embryos were spiked with the corresponding stable-isotope-labelled internal standards (d4-PGE_2_, d4-PGD_2_, d4-6-keto PGF_1α_, d4-PGF_2a_, d4-TxB_2_ and d7-celecoxib). The prostanoid and celecoxib levels were then normalized by total protein concentration.

#### Statistical Analysis

All data were expressed as mean ± S.E.M., unless specified otherwise. Within-group comparison was performed using the two-tailed Student’s t test. All the comparisons were made between animals of the same age and gender on the same genetic background. A *P* value less than 0.05 was considered as a significant difference.

Results of statistical analyses of echo and metabolomics data were corrected for multiple testing within each experiment by the Benjamini-Hochberg method, as implemented by the *p.adjust* function in v4.4.0 of R.

Heatmaps were generated in R using the pheatmap and ggplot2 packages. The abundances of each metabolite were mapped to a common scale in each heatmap as follows: first, the means and SDs of abundances across all samples were calculated separately for each metabolite. Second, the mean abundances across samples within each experimental were calculated separately for each metabolite. Third, these experimental group means were normalized into *Z* scores by subtracting the all-sample means and dividing the result by the all-sample SDs. Again, this procedure was performed separately for each metabolite. For display purposes, any *Z* scores >2 or ≤2 were set to 2 and −2, respectively.

---

**Fig. S1.**
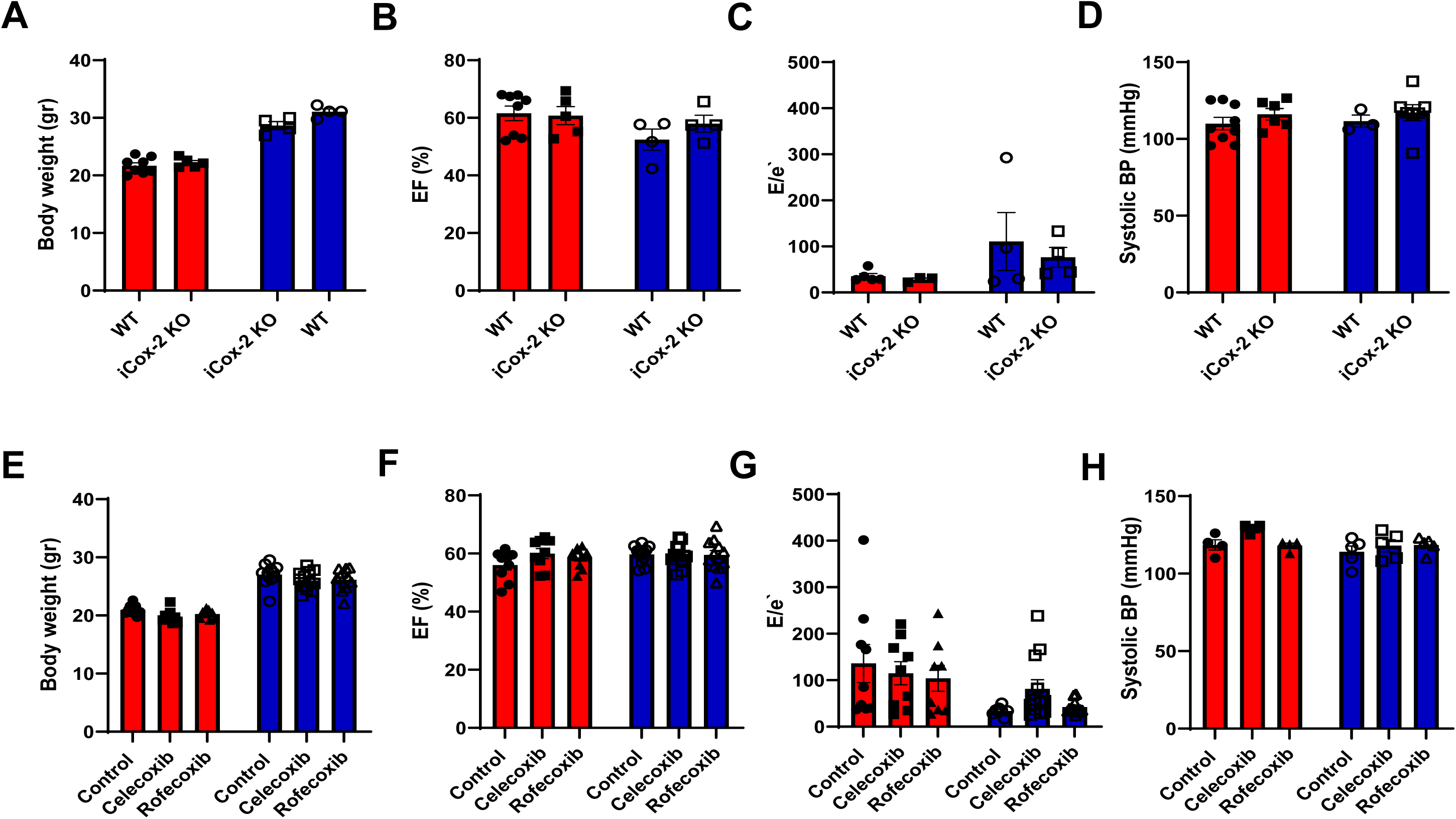
Cox-2 deletion or inhibition does not impair cardiac function in adult male and female mice. (A-D) Cardiac function characterization of 4-5-month-old post-natal tamoxifen inducible Cox-2 KO (iCox-2 KO) female and male mice and their sex- and age- matched littermate controls (WT). (A) Body weight. (B) Ejection fraction (EF). (C) Ratio of peak early Doppler transmitral flow velocity (E) to myocardial tissue Doppler velocity (e′) ratio. (D) Systolic blood pressure (BP) measured by telemetry. (E-H) Cardiac function characterization of 4-5-month-old C57BL/6 female and male mice treated for two months with a control diet, a celecoxib diet (100 mg/Kg) or rofecoxib diet (50 mg/Kg). (E) Body weight. (F) EF (G) E/e′ ratio. (H) Systolic BP measured by telemetry. Pairwise *t* tests were used to determine significant differences between WT and iCox-2 KO within the same sex. Bars (red= females; blue= males) and error bars show means and SEM, respectively, with individual data points superimposed.

**Fig. S2.**
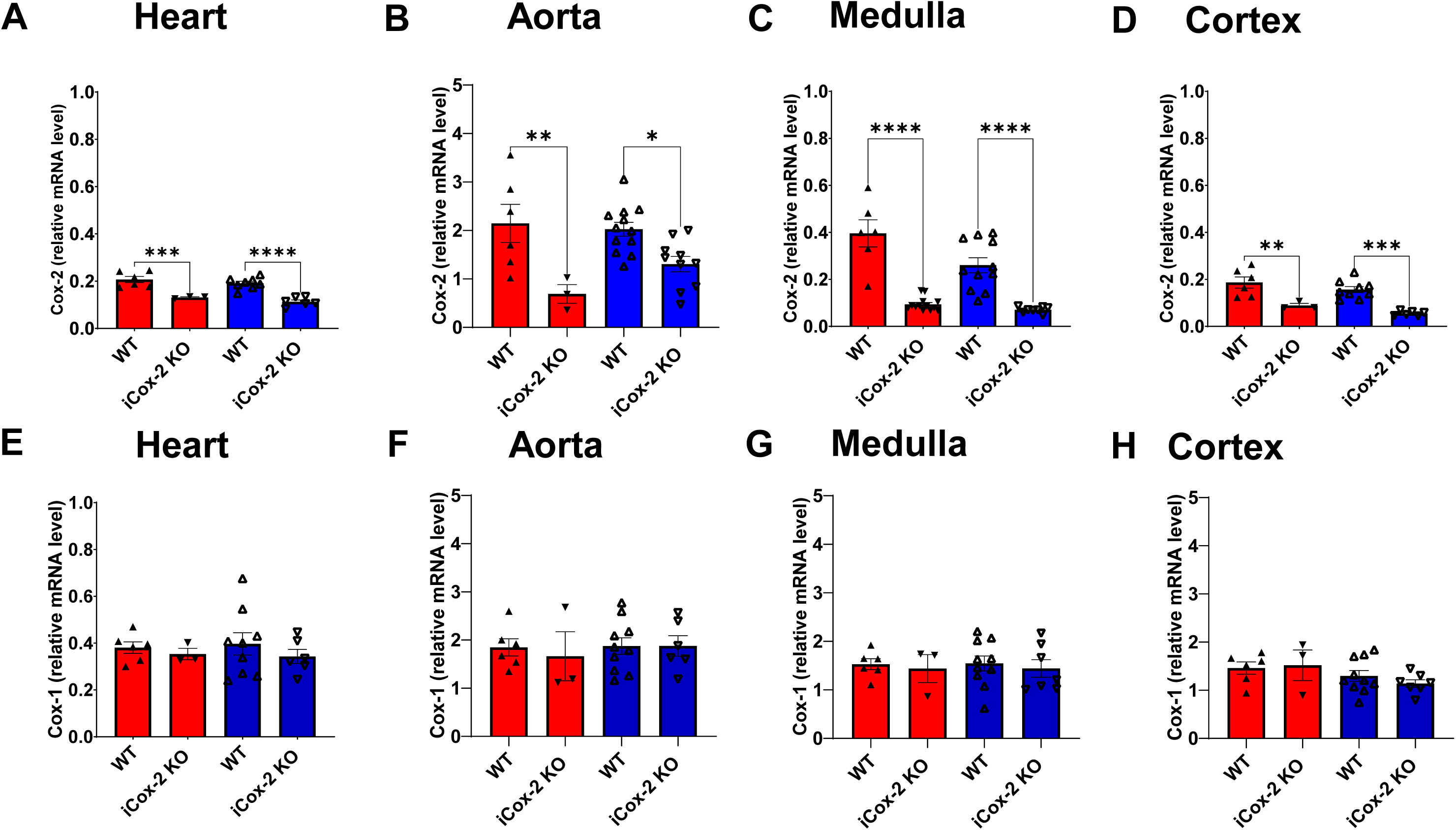
Cox-2 deletion determines a comparable suppression of the COX-2 pathway in aged female and male mice. (A-D) Gene expression of cyclooxygenase (Cox)-2 in aged wild-type (WT) and tamoxifen inducible Cox-2 knockout (iCox-2 KO) female and male mice as detected by RT-PCR. (E-H) Gene expression of Cox-1 in heart, aorta, kidney medulla and kidney cortex of aged wild-type (WT) and tamoxifen inducible Cox-2 knockout (iCox-2 KO) female and male mice as detected by RT-PCR. Pairwise *t* tests were used to determine significant differences between WT and iCox-2 KO within the same sex. Bars (red= females; blue= males) and error bars show means and SEM, respectively, with individual data points superimposed.

**Fig. S3.**
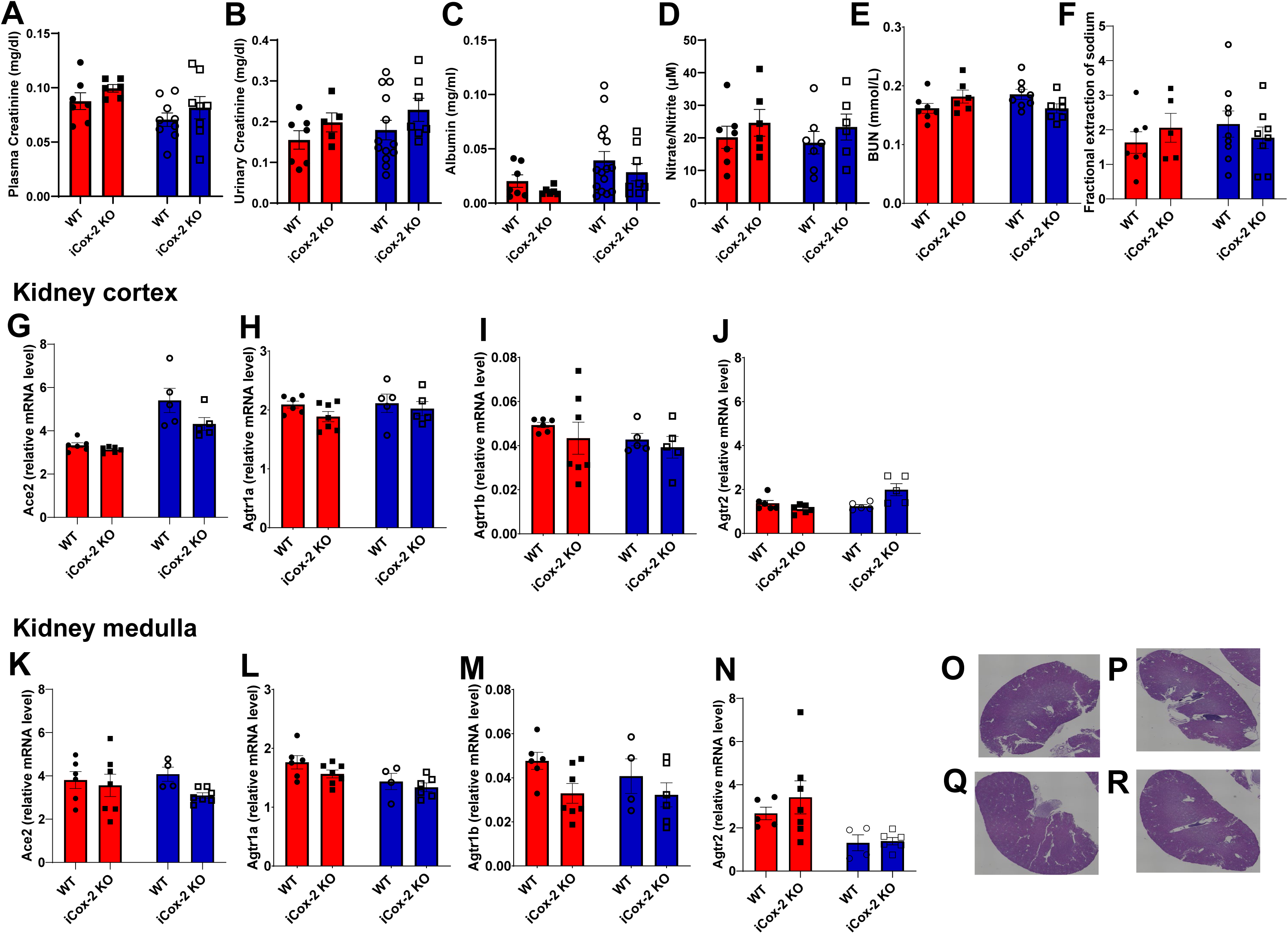
Aged iCox-2 KO female and male mice do not present impairment of kidney function. (A) Plasma creatinine, (B) urinary creatinine, (C) plasma albumin, (D) plasma nitrate/nitrite, (E) blood urea nitrogen (BUN), (F) fractional extraction of sodium measured in aged wild-type (WT) and tamoxifen inducible Cox-2 knockout (iCox-2 KO) female and male mice. (G-J) Gene expression of angiotensin converting enzyme (ace2, G), angiotensin II receptor, type 1a (agtr1a, H), angiotensin II receptor, type 1b (agtr1b, I), and angiotensin II receptor, type 2 (agtr2, J) in the kidney cortex of aged WT and iCox-2 KO female and male mice. (K-N) Gene expression of ace2 (K), agtr1a (L), agtr1b (M), and agtr2 (N) in the kidney medulla of aged WT and iCox-2 KO female and male mice. (O-R) Hematoxylin-eosin (H&E) image of a representative kidney from an aged female WT mouse (O); an aged female iCox-2 KO mouse (P); an aged male WT mouse (Q); an aged male iCox-2 KO mouse (R). The H&E images were taken using light microscopy (Keyence, BZ-X810 microscope) supplied with a digital camera under a magnification of 4 X. Pairwise *t* tests were used to determine significant differences between WT and iCox-2 KO within the same sex. Bars (red= females; blue= males) and error bars show means and SEM, respectively, with individual data points superimposed.

**Fig. S4.**
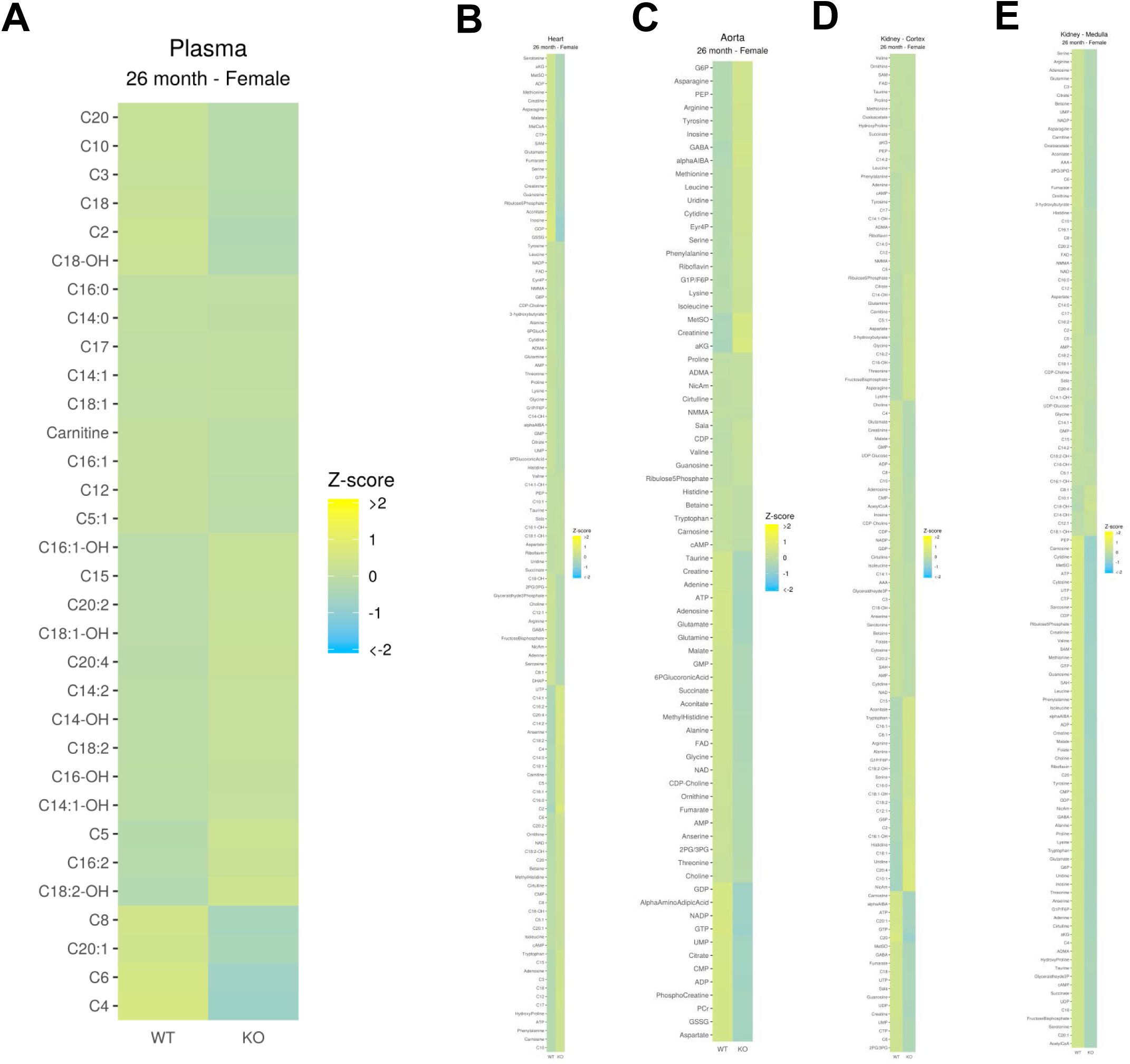
Cox-2 deletion does not impair energy metabolism and oxidative stress in aged female mice. (A) Heatmap (yellow = high, blue = low) showing the relative abundance of acyl- carnitines in plasma from aged WT and iCox-2 KO female mice. (B-E) Heatmap (yellow = high, blue = low) showing the relative abundance of 66 metabolites measured in heart, aorta, renal cortex and renal medulla from aged WT and Cox-2 KO female mice.

**Fig. S5.**
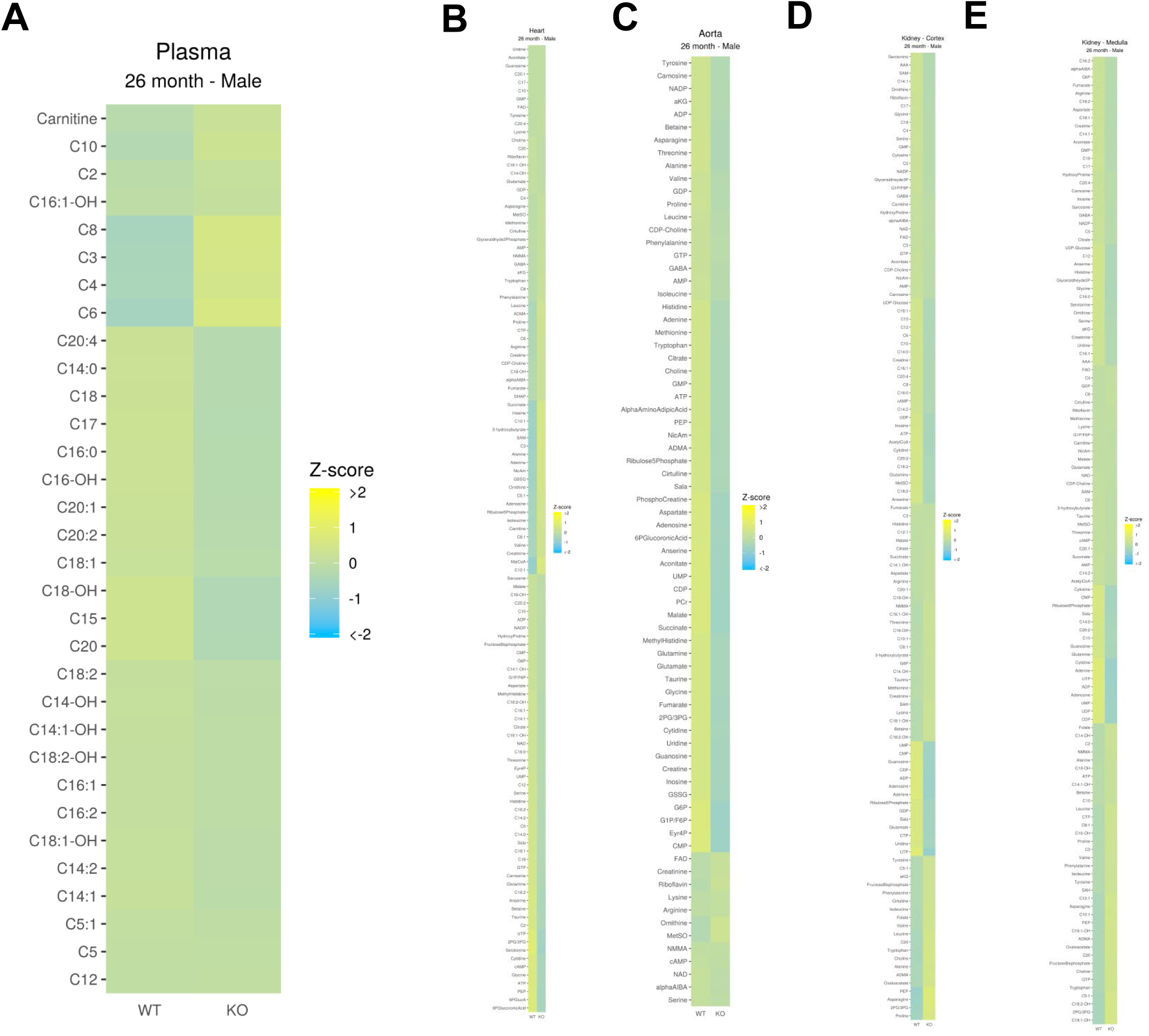
Cox-2 deletion does not impair energy metabolism and oxidative stress in aged male mice. (A) Heatmap ((yellow = high, blue = low) showing the relative abundance of acyl- carnitines in plasma from aged WT and iCox-2 KO male mice. (B-E) Heatmap (yellow = high, blue = low) showing the relative abundance of 66 metabolites measured in heart, aorta, renal cortex and renal medulla from aged WT and Cox-2 KO male mice.

**Fig. S6.**
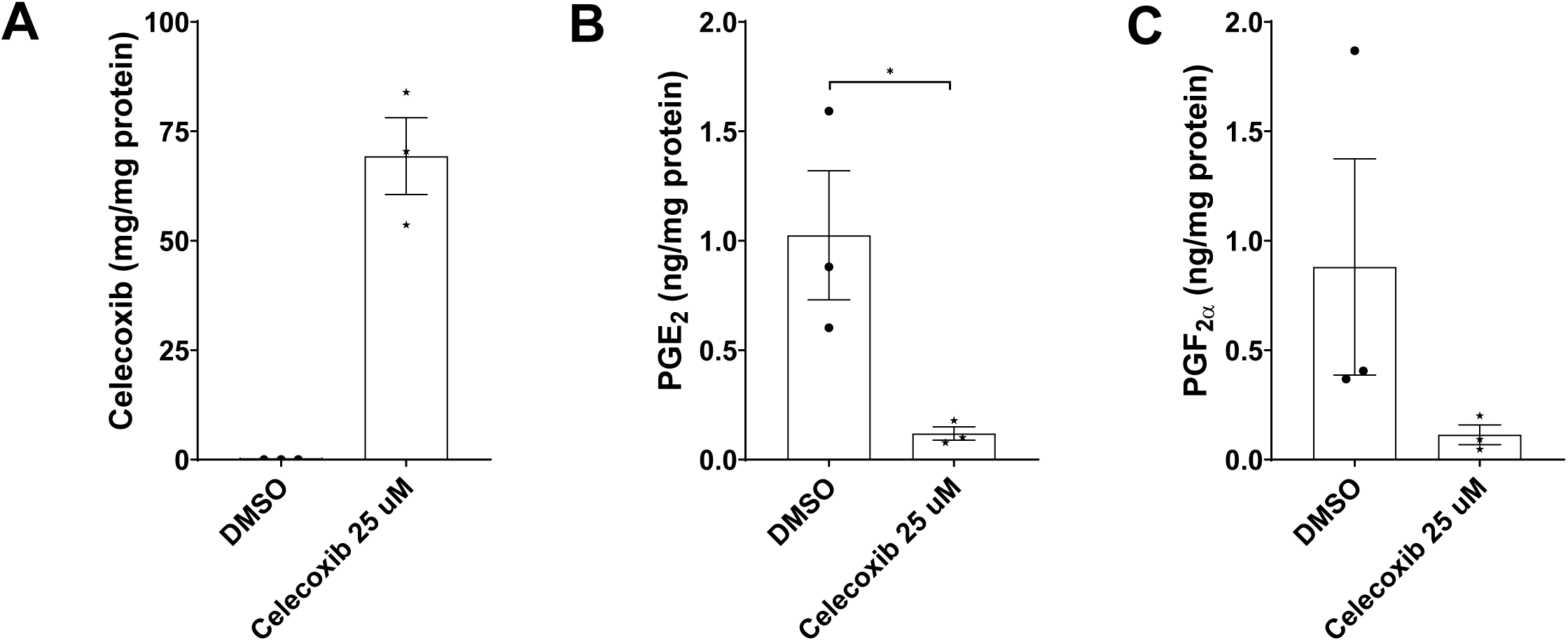
Levels of celecoxib, PGE_2_ and PGF_2α_ in zebrafish homogenizes. (A) intraembryonic celecoxib concentration. (B) intraembryonic PGE_2_ concentration. (C) intraembryonic PGF_2α_ concentration. All measurements were performed using liquid-chromatography mass-spectrometry. Bars and error bars show means and SEM, respectively, with individual data points superimposed.

**Table S1.** Echo parameters in adult female iCox-2 KO and WT mice.

|  | WT (n=5-8) |  |  | iCox-2 KO (n=3-5) |  | p_value |
| --- | --- | --- | --- | --- | --- | --- |
|  | Mean | SD |  | Mean | SD |  |
| A' wave (mm/sec) | 26.10 | 5.40 |  | 26.29 | 9.02 | 0.971 |
| AET (msec) | 47.26 | 5.88 |  | 41.50 | 4.39 | 0.071 |
| CO (ml/min) | 15.08 | 2.06 |  | 16.38 | 3.46 | 0.478 |
| E' wave (mm/sec) | 21.38 | 5.37 |  | 20.34 | 9.51 | 0.850 |
| E'/A' | 0.84 | 0.26 |  | 0.76 | 0.21 | 0.585 |
| EDV (ul) | 54.08 | 6.31 |  | 56.30 | 7.20 | 0.588 |
| ESV (ul) | 20.97 | 5.40 |  | 22.18 | 5.26 | 0.700 |
| FAC (%) | 53.02 | 8.54 |  | 51.24 | 7.37 | 0.699 |
| FS (%) | 31.45 | 4.15 |  | 35.48 | 11.02 | 0.471 |
| HR (bpm) | 457.38 | 52.52 |  | 478.00 | 50.57 | 0.499 |
| IVCT (msec) | 14.07 | 4.38 |  | 12.06 | 3.05 | 0.351 |
| IVRT (msec) | 20.33 | 2.14 |  | 18.72 | 1.97 | 0.200 |
| IVSd (mm) | 0.79 | 0.29 |  | 0.85 | 0.36 | 0.778 |
| IVSs (mm) | 1.10 | 0.33 |  | 1.24 | 0.49 | 0.609 |
| LV mass (mg) | 66.01 | 6.99 |  | 76.46 | 13.50 | 0.166 |
| LV mass/ Body weight | 3.05 | 0.29 |  | 3.45 | 0.65 | 0.246 |
| LVAENDd (mm<sup>2</sup>) | 9.51 | 1.09 |  | 10.08 | 1.38 | 0.464 |
| LVAENDs (mm<sup>2</sup>) | 4.51 | 1.14 |  | 4.94 | 1.14 | 0.525 |
| LVIDd (mm) | 3.36 | 0.19 |  | 3.47 | 0.28 | 0.480 |
| LVIDs (mm) | 2.30 | 0.23 |  | 2.26 | 0.54 | 0.864 |
| LVLd (mm) | 6.83 | 0.36 |  | 6.71 | 0.18 | 0.450 |
| LVLs (mm) | 5.59 | 0.40 |  | 5.38 | 0.16 | 0.228 |
| LVPWd (mm) | 0.71 | 0.14 |  | 0.78 | 0.21 | 0.521 |
| LVPWs (mm) | 0.99 | 0.15 |  | 1.11 | 0.44 | 0.573 |
| MPI | 0.64 | 0.08 |  | 0.65 | 0.07 | 0.865 |
| NFT (msec) | 81.74 | 8.93 |  | 72.72 | 4.97 | 0.039 |
| RWT (%) | 30.61 | 6.51 |  | 31.58 | 8.63 | 0.835 |
| SV (ul) | 33.11 | 4.19 |  | 34.12 | 5.27 | 0.728 |

**Table S2.** Echo parameters in adult male iCox-2 KO and WT mice.

|  | WT (n=4) |  |  | iCox-2 KO (n=4) |  | p_value |
| --- | --- | --- | --- | --- | --- | --- |
|  | Mean | SD |  | Mean | SD |  |
| A' wave (mm/sec) | 25.73 | 6.81 |  | 28.83 | 5.17 | 0.363 |
| AET (msec) | 47.30 | 4.72 |  | 45.90 | 2.29 | 0.484 |
| CO (ml/min) | 20.49 | 3.37 |  | 21.01 | 4.60 | 0.693 |
| E' wave (mm/sec) | 14.19 | 11.36 |  | 12.97 | 5.85 | 0.640 |
| E'/A' | 0.65 | 0.64 |  | 0.47 | 0.24 | 0.468 |
| EDV (ul) | 75.23 | 15.64 |  | 65.98 | 6.65 | 0.431 |
| ESV (ul) | 38.07 | 12.87 |  | 27.70 | 4.73 | 0.312 |
| FAC (%) | 43.14 | 9.34 |  | 48.57 | 7.53 | 0.557 |
| FS (%) | 26.46 | 4.47 |  | 31.07 | 3.18 | 0.235 |
| HR (bpm) | 549.00 | 57.16 |  | 546.25 | 56.94 | 0.613 |
| IVCT (msec) | 12.88 | 5.23 |  | 12.66 | 0.67 | 0.487 |
| IVRT (msec) | 14.15 | 4.10 |  | 17.92 | 1.77 | 0.222 |
| IVSd (mm) | 0.90 | 0.13 |  | 0.88 | 0.11 | 0.406 |
| IVSs (mm) | 1.26 | 0.17 |  | 1.28 | 0.17 | 0.641 |
| LV mass (mg) | 99.52 | 7.31 |  | 85.43 | 11.88 | 0.065 |
| LV mass/ Body weight | 3.25 | 0.18 |  | 2.98 | 0.38 | 0.226 |
| LVAENDd (mm<sup>2</sup>) | 12.56 | 2.12 |  | 11.63 | 0.95 | 0.709 |
| LVAENDs (mm<sup>2</sup>) | 7.29 | 2.32 |  | 5.99 | 1.08 | 0.554 |
| LVAEpid (mm<sup>2</sup>) | 25.49 | 1.97 |  | 23.25 | 2.12 | 0.239 |
| LVIDd (mm) | 3.90 | 0.34 |  | 3.92 | 0.23 | 0.597 |
| LVIDs (mm) | 2.88 | 0.41 |  | 2.70 | 0.22 | 0.783 |
| LVLd (mm) | 7.13 | 0.48 |  | 6.81 | 0.40 | 0.204 |
| LVLs (mm) | 6.21 | 0.26 |  | 5.59 | 0.64 | 0.126 |
| LVPWd (mm) | 0.75 | 0.15 |  | 0.80 | 0.13 | 0.888 |
| LVPWs (mm) | 1.07 | 0.19 |  | 1.22 | 0.15 | 0.472 |
| MPI | 0.57 | 0.12 |  | 0.67 | 0.04 | 0.433 |
| RWT (%) | 29.66 | 4.18 |  | 29.96 | 2.77 | 0.509 |
| SV (ul) | 37.16 | 4.28 |  | 38.28 | 5.98 | 0.884 |

**Table S3.**
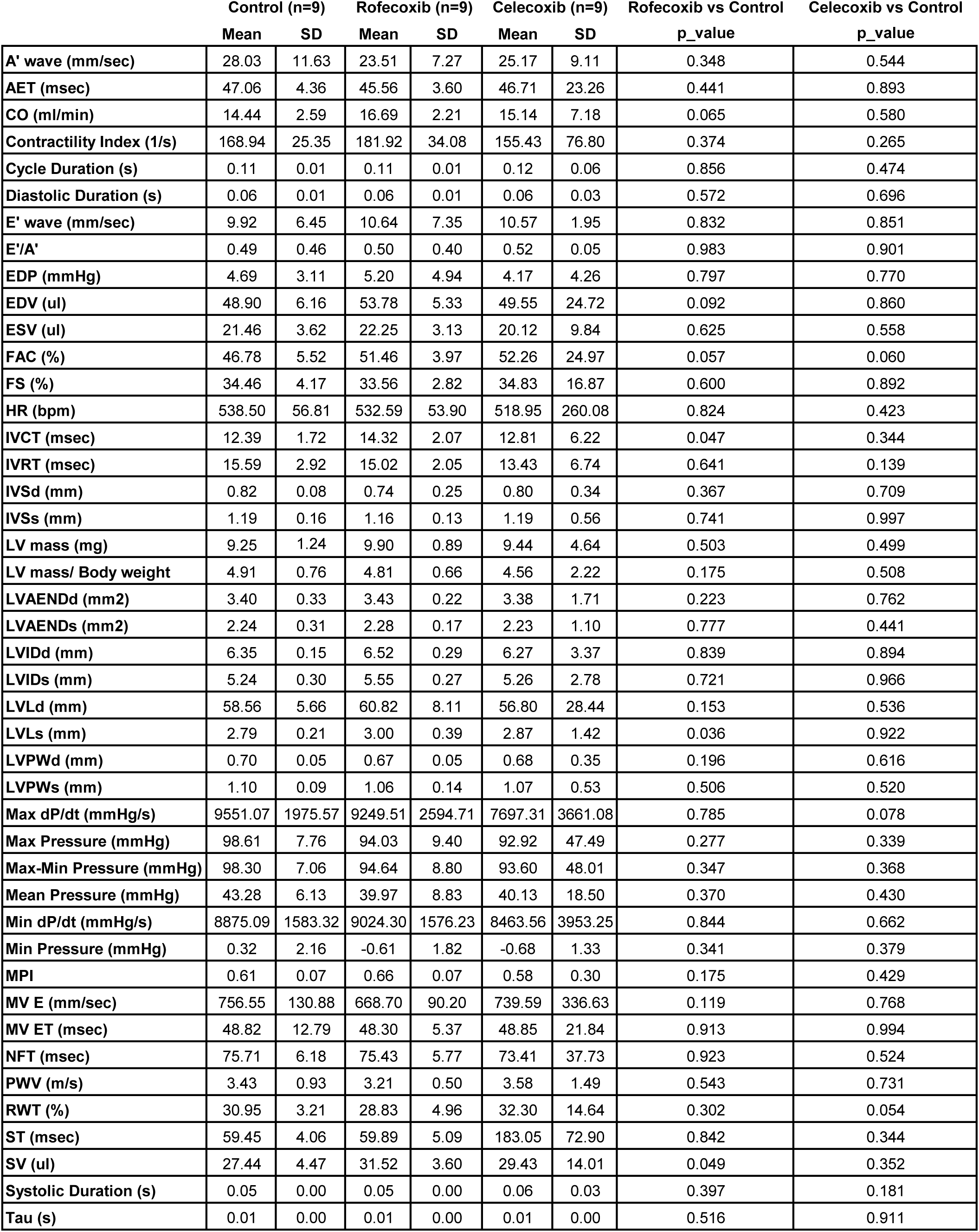
Echo parameters in adult female mice on Celecoxib, Rofecoxib or control diet.

|  | Control (n=9) |  | Rofecoxib (n=9) |  | Celecoxib (n=9) |  | Rofecoxib vs Control | Celecoxib vs Control |
| --- | --- | --- | --- | --- | --- | --- | --- | --- |
|  | Mean | SD | Mean | SD | Mean | SD | p_value | p_value |
| A' wave (mm/sec) | 28.03 | 11.63 | 23.51 | 7.27 | 25.17 | 9.11 | 0.348 | 0.544 |
| AET (msec) | 47.06 | 4.36 | 45.56 | 3.60 | 46.71 | 23.26 | 0.441 | 0.893 |
| CO (ml/min) | 14.44 | 2.59 | 16.69 | 2.21 | 15.14 | 7.18 | 0.065 | 0.580 |
| Contractility Index (1/s) | 168.94 | 25.35 | 181.92 | 34.08 | 155.43 | 76.80 | 0.374 | 0.265 |
| Cycle Duration (s) | 0.11 | 0.01 | 0.11 | 0.01 | 0.12 | 0.06 | 0.856 | 0.474 |
| Diastolic Duration (s) | 0.06 | 0.01 | 0.06 | 0.01 | 0.06 | 0.03 | 0.572 | 0.696 |
| E' wave (mm/sec) | 9.92 | 6.45 | 10.64 | 7.35 | 10.57 | 1.95 | 0.832 | 0.851 |
| E'/A' | 0.49 | 0.46 | 0.50 | 0.40 | 0.52 | 0.05 | 0.983 | 0.901 |
| EDP (mmHg) | 4.69 | 3.11 | 5.20 | 4.94 | 4.17 | 4.26 | 0.797 | 0.770 |
| EDV (ul) | 48.90 | 6.16 | 53.78 | 5.33 | 49.55 | 24.72 | 0.092 | 0.860 |
| ESV (ul) | 21.46 | 3.62 | 22.25 | 3.13 | 20.12 | 9.84 | 0.625 | 0.558 |
| FAC (%) | 46.78 | 5.52 | 51.46 | 3.97 | 52.26 | 24.97 | 0.057 | 0.060 |
| FS (%) | 34.46 | 4.17 | 33.56 | 2.82 | 34.83 | 16.87 | 0.600 | 0.892 |
| HR (bpm) | 538.50 | 56.81 | 532.59 | 53.90 | 518.95 | 260.08 | 0.824 | 0.423 |
| IVCT (msec) | 12.39 | 1.72 | 14.32 | 2.07 | 12.81 | 6.22 | 0.047 | 0.344 |
| IVRT (msec) | 15.59 | 2.92 | 15.02 | 2.05 | 13.43 | 6.74 | 0.641 | 0.139 |
| IVSd (mm) | 0.82 | 0.08 | 0.74 | 0.25 | 0.80 | 0.34 | 0.367 | 0.709 |
| IVSs (mm) | 1.19 | 0.16 | 1.16 | 0.13 | 1.19 | 0.56 | 0.741 | 0.997 |
| LV mass (mg) | 9.25 | 1.24 | 9.90 | 0.89 | 9.44 | 4.64 | 0.503 | 0.499 |
| LV mass/ Body weight | 4.91 | 0.76 | 4.81 | 0.66 | 4.56 | 2.22 | 0.175 | 0.508 |
| LVAENDd (mm2) | 3.40 | 0.33 | 3.43 | 0.22 | 3.38 | 1.71 | 0.223 | 0.762 |
| LVAENDs (mm2) | 2.24 | 0.31 | 2.28 | 0.17 | 2.23 | 1.10 | 0.777 | 0.441 |
| LVIDd (mm) | 6.35 | 0.15 | 6.52 | 0.29 | 6.27 | 3.37 | 0.839 | 0.894 |
| LVIDs (mm) | 5.24 | 0.30 | 5.55 | 0.27 | 5.26 | 2.78 | 0.721 | 0.966 |
| LVLd (mm) | 58.56 | 5.66 | 60.82 | 8.11 | 56.80 | 28.44 | 0.153 | 0.536 |
| LVLs (mm) | 2.79 | 0.21 | 3.00 | 0.39 | 2.87 | 1.42 | 0.036 | 0.922 |
| LVPWd (mm) | 0.70 | 0.05 | 0.67 | 0.05 | 0.68 | 0.35 | 0.196 | 0.616 |
| LVPWs (mm) | 1.10 | 0.09 | 1.06 | 0.14 | 1.07 | 0.53 | 0.506 | 0.520 |
| Max dP/dt (mmHg/s) | 9551.07 | 1975.57 | 9249.51 | 2594.71 | 7697.31 | 3661.08 | 0.785 | 0.078 |
| Max Pressure (mmHg) | 98.61 | 7.76 | 94.03 | 9.40 | 92.92 | 47.49 | 0.277 | 0.339 |
| Max-Min Pressure (mmHg) | 98.30 | 7.06 | 94.64 | 8.80 | 93.60 | 48.01 | 0.347 | 0.368 |
| Mean Pressure (mmHg) | 43.28 | 6.13 | 39.97 | 8.83 | 40.13 | 18.50 | 0.370 | 0.430 |
| Min dP/dt (mmHg/s) | 8875.09 | 1583.32 | 9024.30 | 1576.23 | 8463.56 | 3953.25 | 0.844 | 0.662 |
| Min Pressure (mmHg) | 0.32 | 2.16 | -0.61 | 1.82 | -0.68 | 1.33 | 0.341 | 0.379 |
| MPI | 0.61 | 0.07 | 0.66 | 0.07 | 0.58 | 0.30 | 0.175 | 0.429 |
| MV E (mm/sec) | 756.55 | 130.88 | 668.70 | 90.20 | 739.59 | 336.63 | 0.119 | 0.768 |
| MV ET (msec) | 48.82 | 12.79 | 48.30 | 5.37 | 48.85 | 21.84 | 0.913 | 0.994 |
| NFT (msec) | 75.71 | 6.18 | 75.43 | 5.77 | 73.41 | 37.73 | 0.923 | 0.524 |
| PWV (m/s) | 3.43 | 0.93 | 3.21 | 0.50 | 3.58 | 1.49 | 0.543 | 0.731 |
| RWT (%) | 30.95 | 3.21 | 28.83 | 4.96 | 32.30 | 14.64 | 0.302 | 0.054 |
| ST (msec) | 59.45 | 4.06 | 59.89 | 5.09 | 183.05 | 72.90 | 0.842 | 0.344 |
| SV (ul) | 27.44 | 4.47 | 31.52 | 3.60 | 29.43 | 14.01 | 0.049 | 0.352 |
| Systolic Duration (s) | 0.05 | 0.00 | 0.05 | 0.00 | 0.06 | 0.03 | 0.397 | 0.181 |
| Tau (s) | 0.01 | 0.00 | 0.01 | 0.00 | 0.01 | 0.00 | 0.516 | 0.911 |

**Table S4.**
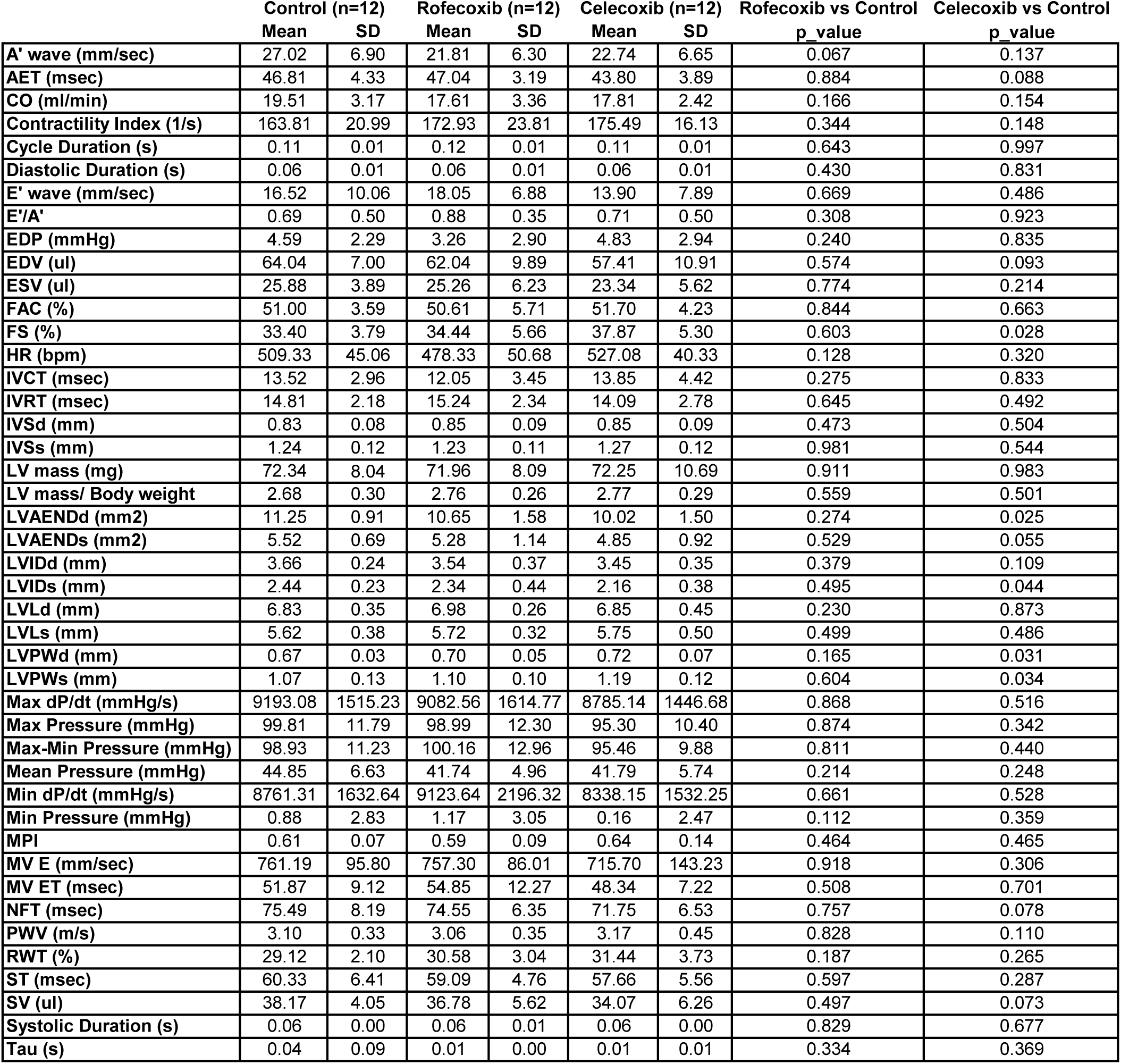
Echo parameters in adult male mice on Celecoxib, Rofecoxib or control diet.

**Table S5.** Echo parameters in old female iCox-2 KO and WT mice.

|  | WT (n=3-6) |  | iCox-2 KO (n=3-4) |  | p_value |
| --- | --- | --- | --- | --- | --- |
|  | Mean | SD | Mean | SD |  |
| A' (mm/s) | 2.39 | 27.88 | 29.92 | 6.28 | 0.061 |
| AET(ms) | 38.42 | 0.97 | 40.65 | 4.89 | 0.516 |
| CO (ml/min) | 20.38 | 2.93 | 17.53 | 2.19 | 0.118 |
| E' (mm/s) | 25.92 | 8.49 | 12.14 | 10.12 | 0.123 |
| E'/A' | 0.11 | 1.16 | 0.40 | 0.33 | 0.354 |
| EAT (ms) | 13.23 | 1.36 | 10.65 | 3.51 | 0.332 |
| FAC (%) | 46.70 | 8.99 | 50.78 | 17.51 | 0.689 |
| FS% | 28.39 | 6.39 | 31.14 | 13.04 | 0.715 |
| HR (bpm) | 553.17 | 22.43 | 536.00 | 29.89 | 0.370 |
| IVCT (ms) | 15.84 | 6.31 | 17.41 | 2.52 | 0.719 |
| IVRT(ms) | 17.59 | 3.93 | 14.81 | 2.92 | 0.385 |
| IVSd(mm) | 1.01 | 0.11 | 0.87 | 0.18 | 0.218 |
| IVSs (mm) | 1.36 | 0.17 | 1.15 | 0.28 | 0.246 |
| Lv mass(mg) | 87.55 | 21.64 | 76.77 | 12.96 | 0.354 |
| LVAENDd(mm2) | 11.60 | 1.87 | 10.80 | 4.39 | 0.750 |
| LVAENDs(mm2) | 6.32 | 2.13 | 5.84 | 4.08 | 0.839 |
| LVEDV(ul) | 66.68 | 13.00 | 63.18 | 29.15 | 0.833 |
| LVESV(ul) | 29.90 | 9.96 | 30.22 | 23.56 | 0.981 |
| LVIDd(mm) | 3.78 | 0.43 | 3.53 | 0.71 | 0.566 |
| LVIDs (mm) | 2.73 | 0.53 | 2.50 | 0.95 | 0.689 |
| LVLd(mm) | 6.87 | 0.46 | 6.92 | 0.40 | 0.875 |
| LVLs(mm) | 5.67 | 0.51 | 5.93 | 0.65 | 0.531 |
| LVPWd(mm) | 1.08 | 0.24 | 0.88 | 0.23 | 0.213 |
| LVPWs(mm) | 1.44 | 0.25 | 1.26 | 0.36 | 0.420 |
| MPI | 0.66 | 0.41 | 0.58 | 0.25 | 0.777 |
| NFT(ms) | 72.04 | 9.15 | 72.96 | 4.10 | 0.884 |
| RWT | 35.72 | 3.02 | 33.40 | 7.84 | 0.606 |
| SV (ul) | 36.78 | 4.50 | 32.96 | 6.23 | 0.338 |

**Table S6.**
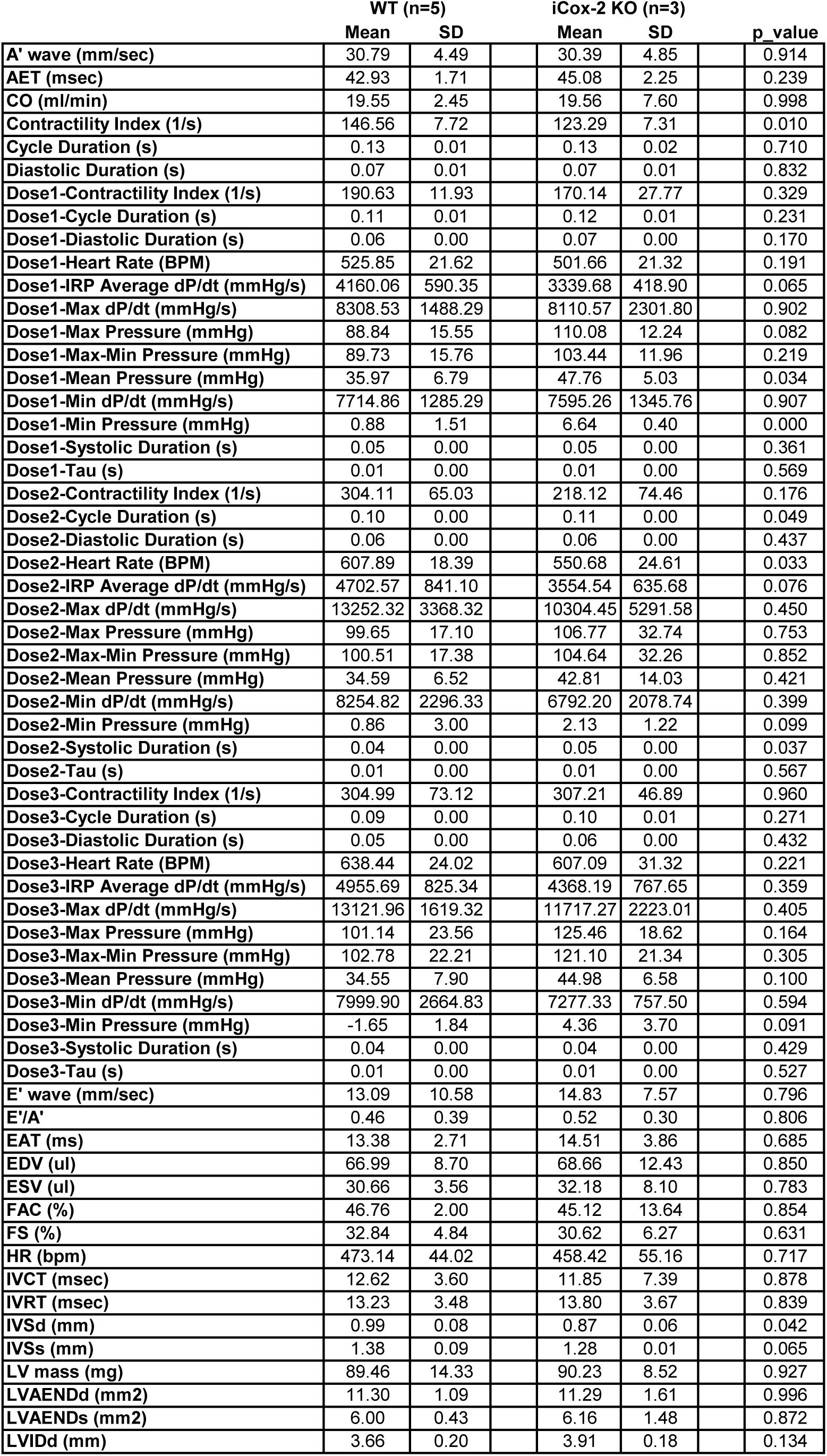
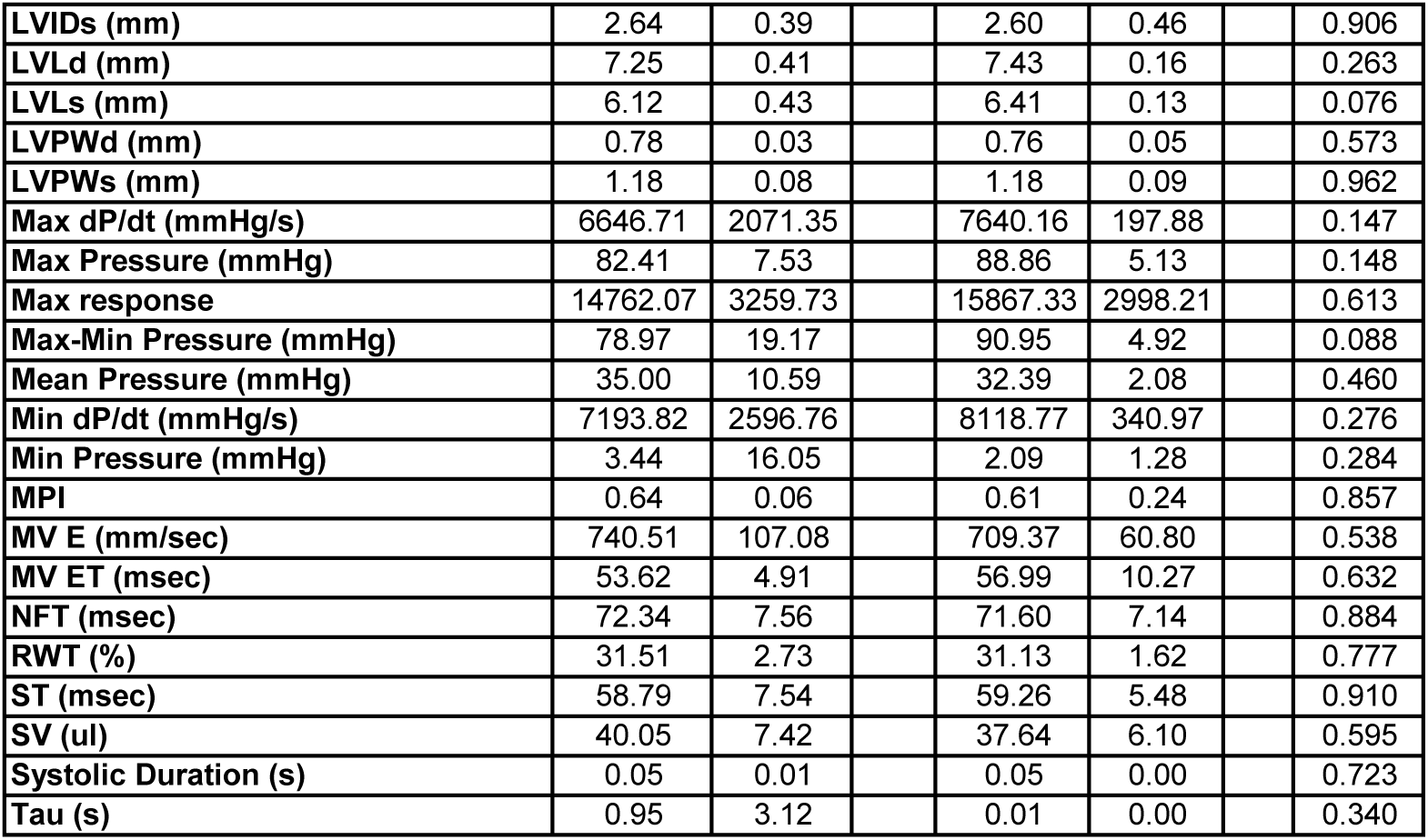
Hemodynamic parameters in old female iCox-2 KO and WT mice.

|  | WT (n=5) |  |  | iCox-2 KO (n=3) |  | p_value |
| --- | --- | --- | --- | --- | --- | --- |
|  | Mean | SD |  | Mean | SD |  |
| A' wave (mm/sec) | 30.79 | 4.49 |  | 30.39 | 4.85 | 0.914 |
| AET (msec) | 42.93 | 1.71 |  | 45.08 | 2.25 | 0.239 |
| CO (ml/min) | 19.55 | 2.45 |  | 19.56 | 7.60 | 0.998 |
| Contractility Index (1/s) | 146.56 | 7.72 |  | 123.29 | 7.31 | 0.010 |
| Cycle Duration (s) | 0.13 | 0.01 |  | 0.13 | 0.02 | 0.710 |
| Diastolic Duration (s) | 0.07 | 0.01 |  | 0.07 | 0.01 | 0.832 |
| Dose1-Contractility Index (1/s) | 190.63 | 11.93 |  | 170.14 | 27.77 | 0.329 |
| Dose1-Cycle Duration (s) | 0.11 | 0.01 |  | 0.12 | 0.01 | 0.231 |
| Dose1-Diastolic Duration (s) | 0.06 | 0.00 |  | 0.07 | 0.00 | 0.170 |
| Dose1-Heart Rate (BPM) | 525.85 | 21.62 |  | 501.66 | 21.32 | 0.191 |
| Dose1-IRP Average dP/dt (mmHg/s) | 4160.06 | 590.35 |  | 3339.68 | 418.90 | 0.065 |
| Dose1-Max dP/dt (mmHg/s) | 8308.53 | 1488.29 |  | 8110.57 | 2301.80 | 0.902 |
| Dose1-Max Pressure (mmHg) | 88.84 | 15.55 |  | 110.08 | 12.24 | 0.082 |
| Dose1-Max-Min Pressure (mmHg) | 89.73 | 15.76 |  | 103.44 | 11.96 | 0.219 |
| Dose1-Mean Pressure (mmHg) | 35.97 | 6.79 |  | 47.76 | 5.03 | 0.034 |
| Dose1-Min dP/dt (mmHg/s) | 7714.86 | 1285.29 |  | 7595.26 | 1345.76 | 0.907 |
| Dose1-Min Pressure (mmHg) | 0.88 | 1.51 |  | 6.64 | 0.40 | 0.000 |
| Dose1-Systolic Duration (s) | 0.05 | 0.00 |  | 0.05 | 0.00 | 0.361 |
| Dose1-Tau (s) | 0.01 | 0.00 |  | 0.01 | 0.00 | 0.569 |
| Dose2-Contractility Index (1/s) | 304.11 | 65.03 |  | 218.12 | 74.46 | 0.176 |
| Dose2-Cycle Duration (s) | 0.10 | 0.00 |  | 0.11 | 0.00 | 0.049 |
| Dose2-Diastolic Duration (s) | 0.06 | 0.00 |  | 0.06 | 0.00 | 0.437 |
| Dose2-Heart Rate (BPM) | 607.89 | 18.39 |  | 550.68 | 24.61 | 0.033 |
| Dose2-IRP Average dP/dt (mmHg/s) | 4702.57 | 841.10 |  | 3554.54 | 635.68 | 0.076 |
| Dose2-Max dP/dt (mmHg/s) | 13252.32 | 3368.32 |  | 10304.45 | 5291.58 | 0.450 |
| Dose2-Max Pressure (mmHg) | 99.65 | 17.10 |  | 106.77 | 32.74 | 0.753 |
| Dose2-Max-Min Pressure (mmHg) | 100.51 | 17.38 |  | 104.64 | 32.26 | 0.852 |
| Dose2-Mean Pressure (mmHg) | 34.59 | 6.52 |  | 42.81 | 14.03 | 0.421 |
| Dose2-Min dP/dt (mmHg/s) | 8254.82 | 2296.33 |  | 6792.20 | 2078.74 | 0.399 |
| Dose2-Min Pressure (mmHg) | 0.86 | 3.00 |  | 2.13 | 1.22 | 0.099 |
| Dose2-Systolic Duration (s) | 0.04 | 0.00 |  | 0.05 | 0.00 | 0.037 |
| Dose2-Tau (s) | 0.01 | 0.00 |  | 0.01 | 0.00 | 0.567 |
| Dose3-Contractility Index (1/s) | 304.99 | 73.12 |  | 307.21 | 46.89 | 0.960 |
| Dose3-Cycle Duration (s) | 0.09 | 0.00 |  | 0.10 | 0.01 | 0.271 |
| Dose3-Diastolic Duration (s) | 0.05 | 0.00 |  | 0.06 | 0.00 | 0.432 |
| Dose3-Heart Rate (BPM) | 638.44 | 24.02 |  | 607.09 | 31.32 | 0.221 |
| Dose3-IRP Average dP/dt (mmHg/s) | 4955.69 | 825.34 |  | 4368.19 | 767.65 | 0.359 |
| Dose3-Max dP/dt (mmHg/s) | 13121.96 | 1619.32 |  | 11717.27 | 2223.01 | 0.405 |
| Dose3-Max Pressure (mmHg) | 101.14 | 23.56 |  | 125.46 | 18.62 | 0.164 |
| Dose3-Max-Min Pressure (mmHg) | 102.78 | 22.21 |  | 121.10 | 21.34 | 0.305 |
| Dose3-Mean Pressure (mmHg) | 34.55 | 7.90 |  | 44.98 | 6.58 | 0.100 |
| Dose3-Min dP/dt (mmHg/s) | 7999.90 | 2664.83 |  | 7277.33 | 757.50 | 0.594 |
| Dose3-Min Pressure (mmHg) | -1.65 | 1.84 |  | 4.36 | 3.70 | 0.091 |
| Dose3-Systolic Duration (s) | 0.04 | 0.00 |  | 0.04 | 0.00 | 0.429 |
| Dose3-Tau (s) | 0.01 | 0.00 |  | 0.01 | 0.00 | 0.527 |
| E' wave (mm/sec) | 13.09 | 10.58 |  | 14.83 | 7.57 | 0.796 |
| E'/A' | 0.46 | 0.39 |  | 0.52 | 0.30 | 0.806 |
| EAT (ms) | 13.38 | 2.71 |  | 14.51 | 3.86 | 0.685 |
| EDV (ul) | 66.99 | 8.70 |  | 68.66 | 12.43 | 0.850 |
| ESV (ul) | 30.66 | 3.56 |  | 32.18 | 8.10 | 0.783 |
| FAC (%) | 46.76 | 2.00 |  | 45.12 | 13.64 | 0.854 |
| FS (%) | 32.84 | 4.84 |  | 30.62 | 6.27 | 0.631 |
| HR (bpm) | 473.14 | 44.02 |  | 458.42 | 55.16 | 0.717 |
| IVCT (msec) | 12.62 | 3.60 |  | 11.85 | 7.39 | 0.878 |
| IVRT (msec) | 13.23 | 3.48 |  | 13.80 | 3.67 | 0.839 |
| IVSd (mm) | 0.99 | 0.08 |  | 0.87 | 0.06 | 0.042 |
| IVSs (mm) | 1.38 | 0.09 |  | 1.28 | 0.01 | 0.065 |
| LV mass (mg) | 89.46 | 14.33 |  | 90.23 | 8.52 | 0.927 |
| LVAENDd (mm2) | 11.30 | 1.09 |  | 11.29 | 1.61 | 0.996 |
| LVAENDs (mm2) | 6.00 | 0.43 |  | 6.16 | 1.48 | 0.872 |
| LVIDd (mm) | 3.66 | 0.20 |  | 3.91 | 0.18 | 0.134 |

|  |  |  |  |  |  |  |  |
| --- | --- | --- | --- | --- | --- | --- | --- |
| **LVIDs (mm)** | 2.64 | 0.39 |  | 2.60 | 0.46 |  | 0.906 |
| **LVLd (mm)** | 7.25 | 0.41 |  | 7.43 | 0.16 |  | 0.263 |
| **LVLs (mm)** | 6.12 | 0.43 |  | 6.41 | 0.13 |  | 0.076 |
| **LVPWd (mm)** | 0.78 | 0.03 |  | 0.76 | 0.05 |  | 0.573 |
| **LVPWs (mm)** | 1.18 | 0.08 |  | 1.18 | 0.09 |  | 0.962 |
| **Max dP/dt (mmHg/s)** | 6646.71 | 2071.35 |  | 7640.16 | 197.88 |  | 0.147 |
| **Max Pressure (mmHg)** | 82.41 | 7.53 |  | 88.86 | 5.13 |  | 0.148 |
| **Max response** | 14762.07 | 3259.73 |  | 15867.33 | 2998.21 |  | 0.613 |
| **Max-Min Pressure (mmHg)** | 78.97 | 19.17 |  | 90.95 | 4.92 |  | 0.088 |
| **Mean Pressure (mmHg)** | 35.00 | 10.59 |  | 32.39 | 2.08 |  | 0.460 |
| **Min dP/dt (mmHg/s)** | 7193.82 | 2596.76 |  | 8118.77 | 340.97 |  | 0.276 |
| **Min Pressure (mmHg)** | 3.44 | 16.05 |  | 2.09 | 1.28 |  | 0.284 |
| **MPI** | 0.64 | 0.06 |  | 0.61 | 0.24 |  | 0.857 |
| **MV E (mm/sec)** | 740.51 | 107.08 |  | 709.37 | 60.80 |  | 0.538 |
| **MV ET (msec)** | 53.62 | 4.91 |  | 56.99 | 10.27 |  | 0.632 |
| **NFT (msec)** | 72.34 | 7.56 |  | 71.60 | 7.14 |  | 0.884 |
| **RWT (%)** | 31.51 | 2.73 |  | 31.13 | 1.62 |  | 0.777 |
| **ST (msec)** | 58.79 | 7.54 |  | 59.26 | 5.48 |  | 0.910 |
| **SV (ul)** | 40.05 | 7.42 |  | 37.64 | 6.10 |  | 0.595 |
| **Systolic Duration (s)** | 0.05 | 0.01 |  | 0.05 | 0.00 |  | 0.723 |
| **Tau (s)** | 0.95 | 3.12 |  | 0.01 | 0.00 |  | 0.340 |

**Table S7.** Echo parameters in old male iCox-2 KO and WT mice.

|  | WT (n=3-4) |  |  | iCox-2 KO (n=3-4) |  | p_value |
| --- | --- | --- | --- | --- | --- | --- |
|  | Mean | SD |  | Mean | SD |  |
| A' (mm/s) | 23.283 | 7.952 |  | 20.131 | 11.121 | 0.663 |
| AET (ms) | 46.250 | 5.119 |  | 47.915 | 8.906 | 0.759 |
| CO (ml/min) | 17.460 | 2.685 |  | 21.285 | 6.074 | 0.394 |
| E' (mm/s) | 22.930 | 7.062 |  | 20.313 | 3.852 | 0.546 |
| E'/A' | 1.015 | 0.251 |  | 1.652 | 1.581 | 0.482 |
| EAT (ms) | 13.128 | 1.717 |  | 16.668 | 5.859 | 0.319 |
| FAC (%) | 40.820 | 4.403 |  | 69.445 | 16.994 | 0.094 |
| FS% | 26.990 | 6.625 |  | 40.213 | 18.171 | 0.247 |
| HR (bpm) | 455.500 | 38.179 |  | 499.750 | 74.173 | 0.342 |
| IVCT (ms) | 16.320 | 3.364 |  | 21.598 | 16.899 | 0.581 |
| IVRT(ms) | 18.193 | 6.153 |  | 12.640 | 2.709 | 0.172 |
| IVSd(mm) | 0.791 | 0.101 |  | 0.860 | 0.225 | 0.606 |
| IVSs (mm) | 1.002 | 0.129 |  | 1.211 | 0.360 | 0.339 |
| Lv mass(mg) | 84.406 | 15.983 |  | 72.170 | 24.916 | 0.508 |
| LVAENDd(mm2) | 12.678 | 2.062 |  | 9.578 | 4.114 | 0.324 |
| LVAENDs(mm2) | 7.555 | 1.663 |  | 3.392 | 3.164 | 0.135 |
| LVEDV(ul) | 78.340 | 15.933 |  | 61.607 | 30.739 | 0.457 |
| LVESV(ul) | 40.124 | 12.644 |  | 18.053 | 20.142 | 0.189 |
| LVIDd(mm) | 4.167 | 0.266 |  | 3.619 | 0.931 | 0.329 |
| LVIDs (mm) | 3.055 | 0.472 |  | 2.286 | 1.179 | 0.293 |
| LVLd(mm) | 7.376 | 0.443 |  | 7.480 | 0.452 | 0.754 |
| LVLs(mm) | 6.259 | 0.716 |  | 5.832 | 1.110 | 0.546 |
| LVPWd(mm) | 0.839 | 0.101 |  | 0.783 | 0.044 | 0.358 |
| LVPWs(mm) | 1.191 | 0.139 |  | 1.269 | 0.259 | 0.620 |
| MPI | 0.428 | 0.049 |  | 0.405 | 0.085 | 0.656 |
| NFT(ms) | 80.903 | 5.838 |  | 82.083 | 21.685 | 0.922 |
| RWT | 28.090 | 2.320 |  | 32.000 | 7.387 | 0.376 |
| SV (ul) | 38.216 | 3.910 |  | 43.554 | 11.215 | 0.501 |

**Table S8.**
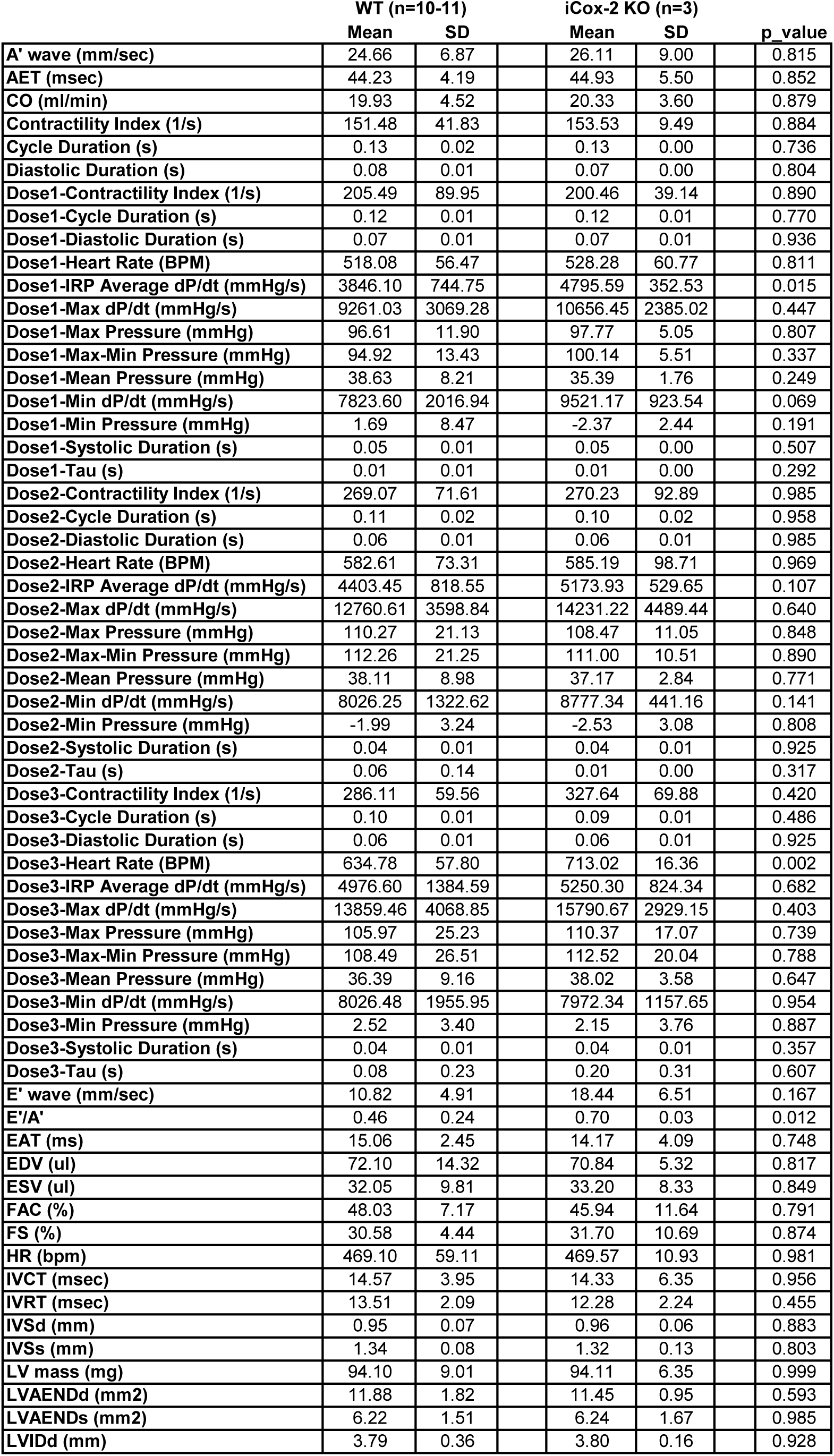
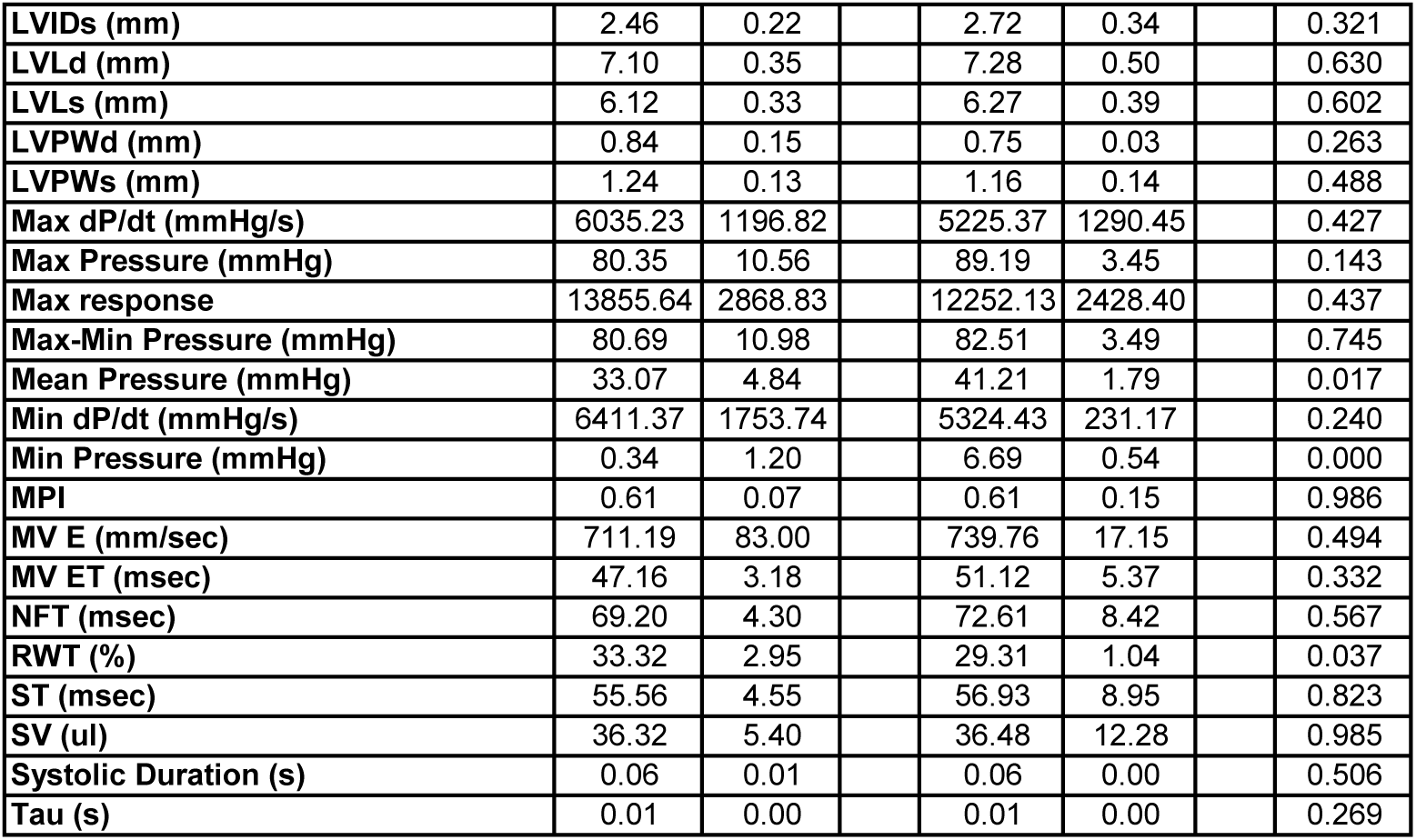
Invasive hemodynamic parameters in old male iCox-2 KO and WT mice.

|  | WT (n=10-11) |  |  | iCox-2 KO (n=3) |  | p_value |
| --- | --- | --- | --- | --- | --- | --- |
|  | Mean | SD |  | Mean | SD |  |
| A' wave (mm/sec) | 24.66 | 6.87 |  | 26.11 | 9.00 | 0.815 |
| AET (msec) | 44.23 | 4.19 |  | 44.93 | 5.50 | 0.852 |
| CO (ml/min) | 19.93 | 4.52 |  | 20.33 | 3.60 | 0.879 |
| Contractility Index (1/s) | 151.48 | 41.83 |  | 153.53 | 9.49 | 0.884 |
| Cycle Duration (s) | 0.13 | 0.02 |  | 0.13 | 0.00 | 0.736 |
| Diastolic Duration (s) | 0.08 | 0.01 |  | 0.07 | 0.00 | 0.804 |
| Dose1-Contractility Index (1/s) | 205.49 | 89.95 |  | 200.46 | 39.14 | 0.890 |
| Dose1-Cycle Duration (s) | 0.12 | 0.01 |  | 0.12 | 0.01 | 0.770 |
| Dose1-Diastolic Duration (s) | 0.07 | 0.01 |  | 0.07 | 0.01 | 0.936 |
| Dose1-Heart Rate (BPM) | 518.08 | 56.47 |  | 528.28 | 60.77 | 0.811 |
| Dose1-IRP Average dP/dt (mmHg/s) | 3846.10 | 744.75 |  | 4795.59 | 352.53 | 0.015 |
| Dose1-Max dP/dt (mmHg/s) | 9261.03 | 3069.28 |  | 10656.45 | 2385.02 | 0.447 |
| Dose1-Max Pressure (mmHg) | 96.61 | 11.90 |  | 97.77 | 5.05 | 0.807 |
| Dose1-Max-Min Pressure (mmHg) | 94.92 | 13.43 |  | 100.14 | 5.51 | 0.337 |
| Dose1-Mean Pressure (mmHg) | 38.63 | 8.21 |  | 35.39 | 1.76 | 0.249 |
| Dose1-Min dP/dt (mmHg/s) | 7823.60 | 2016.94 |  | 9521.17 | 923.54 | 0.069 |
| Dose1-Min Pressure (mmHg) | 1.69 | 8.47 |  | -2.37 | 2.44 | 0.191 |
| Dose1-Systolic Duration (s) | 0.05 | 0.01 |  | 0.05 | 0.00 | 0.507 |
| Dose1-Tau (s) | 0.01 | 0.01 |  | 0.01 | 0.00 | 0.292 |
| Dose2-Contractility Index (1/s) | 269.07 | 71.61 |  | 270.23 | 92.89 | 0.985 |
| Dose2-Cycle Duration (s) | 0.11 | 0.02 |  | 0.10 | 0.02 | 0.958 |
| Dose2-Diastolic Duration (s) | 0.06 | 0.01 |  | 0.06 | 0.01 | 0.985 |
| Dose2-Heart Rate (BPM) | 582.61 | 73.31 |  | 585.19 | 98.71 | 0.969 |
| Dose2-IRP Average dP/dt (mmHg/s) | 4403.45 | 818.55 |  | 5173.93 | 529.65 | 0.107 |
| Dose2-Max dP/dt (mmHg/s) | 12760.61 | 3598.84 |  | 14231.22 | 4489.44 | 0.640 |
| Dose2-Max Pressure (mmHg) | 110.27 | 21.13 |  | 108.47 | 11.05 | 0.848 |
| Dose2-Max-Min Pressure (mmHg) | 112.26 | 21.25 |  | 111.00 | 10.51 | 0.890 |
| Dose2-Mean Pressure (mmHg) | 38.11 | 8.98 |  | 37.17 | 2.84 | 0.771 |
| Dose2-Min dP/dt (mmHg/s) | 8026.25 | 1322.62 |  | 8777.34 | 441.16 | 0.141 |
| Dose2-Min Pressure (mmHg) | -1.99 | 3.24 |  | -2.53 | 3.08 | 0.808 |
| Dose2-Systolic Duration (s) | 0.04 | 0.01 |  | 0.04 | 0.01 | 0.925 |
| Dose2-Tau (s) | 0.06 | 0.14 |  | 0.01 | 0.00 | 0.317 |
| Dose3-Contractility Index (1/s) | 286.11 | 59.56 |  | 327.64 | 69.88 | 0.420 |
| Dose3-Cycle Duration (s) | 0.10 | 0.01 |  | 0.09 | 0.01 | 0.486 |
| Dose3-Diastolic Duration (s) | 0.06 | 0.01 |  | 0.06 | 0.01 | 0.925 |
| Dose3-Heart Rate (BPM) | 634.78 | 57.80 |  | 713.02 | 16.36 | 0.002 |
| Dose3-IRP Average dP/dt (mmHg/s) | 4976.60 | 1384.59 |  | 5250.30 | 824.34 | 0.682 |
| Dose3-Max dP/dt (mmHg/s) | 13859.46 | 4068.85 |  | 15790.67 | 2929.15 | 0.403 |
| Dose3-Max Pressure (mmHg) | 105.97 | 25.23 |  | 110.37 | 17.07 | 0.739 |
| Dose3-Max-Min Pressure (mmHg) | 108.49 | 26.51 |  | 112.52 | 20.04 | 0.788 |
| Dose3-Mean Pressure (mmHg) | 36.39 | 9.16 |  | 38.02 | 3.58 | 0.647 |
| Dose3-Min dP/dt (mmHg/s) | 8026.48 | 1955.95 |  | 7972.34 | 1157.65 | 0.954 |
| Dose3-Min Pressure (mmHg) | 2.52 | 3.40 |  | 2.15 | 3.76 | 0.887 |
| Dose3-Systolic Duration (s) | 0.04 | 0.01 |  | 0.04 | 0.01 | 0.357 |
| Dose3-Tau (s) | 0.08 | 0.23 |  | 0.20 | 0.31 | 0.607 |
| E' wave (mm/sec) | 10.82 | 4.91 |  | 18.44 | 6.51 | 0.167 |
| E'/A' | 0.46 | 0.24 |  | 0.70 | 0.03 | 0.012 |
| EAT (ms) | 15.06 | 2.45 |  | 14.17 | 4.09 | 0.748 |
| EDV (ul) | 72.10 | 14.32 |  | 70.84 | 5.32 | 0.817 |
| ESV (ul) | 32.05 | 9.81 |  | 33.20 | 8.33 | 0.849 |
| FAC (%) | 48.03 | 7.17 |  | 45.94 | 11.64 | 0.791 |
| FS (%) | 30.58 | 4.44 |  | 31.70 | 10.69 | 0.874 |
| HR (bpm) | 469.10 | 59.11 |  | 469.57 | 10.93 | 0.981 |
| IVCT (msec) | 14.57 | 3.95 |  | 14.33 | 6.35 | 0.956 |
| IVRT (msec) | 13.51 | 2.09 |  | 12.28 | 2.24 | 0.455 |
| IVSd (mm) | 0.95 | 0.07 |  | 0.96 | 0.06 | 0.883 |
| IVSs (mm) | 1.34 | 0.08 |  | 1.32 | 0.13 | 0.803 |
| LV mass (mg) | 94.10 | 9.01 |  | 94.11 | 6.35 | 0.999 |
| LVAENDd (mm2) | 11.88 | 1.82 |  | 11.45 | 0.95 | 0.593 |
| LVAENDs (mm2) | 6.22 | 1.51 |  | 6.24 | 1.67 | 0.985 |
| LVIDd (mm) | 3.79 | 0.36 |  | 3.80 | 0.16 | 0.928 |

|  |  |  |  |  |  |  |  |
| --- | --- | --- | --- | --- | --- | --- | --- |
| **LVIDs (mm)** | 2.46 | 0.22 |  | 2.72 | 0.34 |  | 0.321 |
| **LVLd (mm)** | 7.10 | 0.35 |  | 7.28 | 0.50 |  | 0.630 |
| **LVLs (mm)** | 6.12 | 0.33 |  | 6.27 | 0.39 |  | 0.602 |
| **LVPWd (mm)** | 0.84 | 0.15 |  | 0.75 | 0.03 |  | 0.263 |
| **LVPWs (mm)** | 1.24 | 0.13 |  | 1.16 | 0.14 |  | 0.488 |
| **Max dP/dt (mmHg/s)** | 6035.23 | 1196.82 |  | 5225.37 | 1290.45 |  | 0.427 |
| **Max Pressure (mmHg)** | 80.35 | 10.56 |  | 89.19 | 3.45 |  | 0.143 |
| **Max response** | 13855.64 | 2868.83 |  | 12252.13 | 2428.40 |  | 0.437 |
| **Max-Min Pressure (mmHg)** | 80.69 | 10.98 |  | 82.51 | 3.49 |  | 0.745 |
| **Mean Pressure (mmHg)** | 33.07 | 4.84 |  | 41.21 | 1.79 |  | 0.017 |
| **Min dP/dt (mmHg/s)** | 6411.37 | 1753.74 |  | 5324.43 | 231.17 |  | 0.240 |
| **Min Pressure (mmHg)** | 0.34 | 1.20 |  | 6.69 | 0.54 |  | 0.000 |
| **MPI** | 0.61 | 0.07 |  | 0.61 | 0.15 |  | 0.986 |
| **MV E (mm/sec)** | 711.19 | 83.00 |  | 739.76 | 17.15 |  | 0.494 |
| **MV ET (msec)** | 47.16 | 3.18 |  | 51.12 | 5.37 |  | 0.332 |
| **NFT (msec)** | 69.20 | 4.30 |  | 72.61 | 8.42 |  | 0.567 |
| **RWT (%)** | 33.32 | 2.95 |  | 29.31 | 1.04 |  | 0.037 |
| **ST (msec)** | 55.56 | 4.55 |  | 56.93 | 8.95 |  | 0.823 |
| **SV (ul)** | 36.32 | 5.40 |  | 36.48 | 12.28 |  | 0.985 |
| **Systolic Duration (s)** | 0.06 | 0.01 |  | 0.06 | 0.00 |  | 0.506 |
| **Tau (s)** | 0.01 | 0.00 |  | 0.01 | 0.00 |  | 0.269 |

**Table S9.**
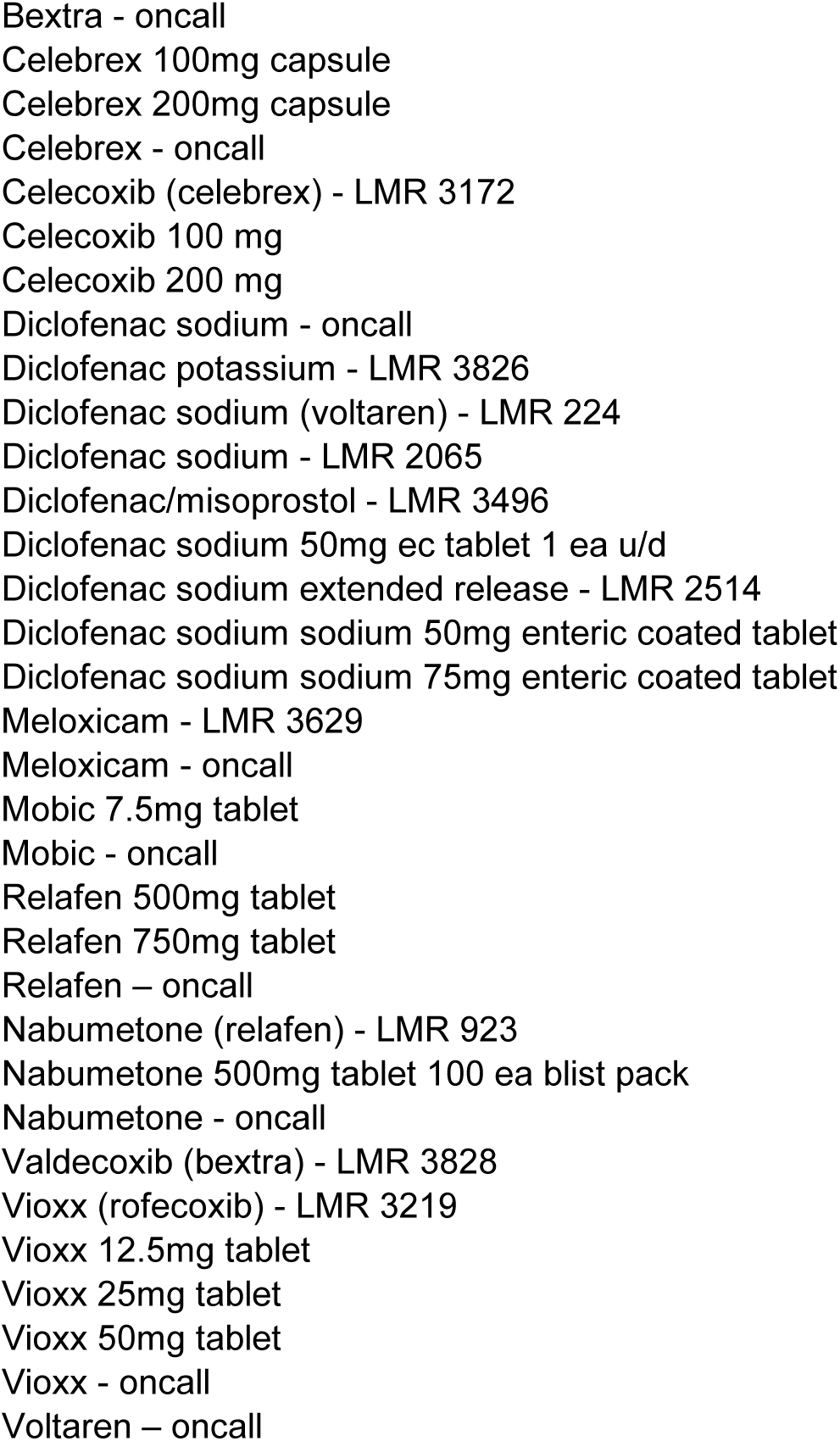
NSAIDs selective for COX-2 inhibition prescriptions extracted from the EMR database.

**Table S10.**
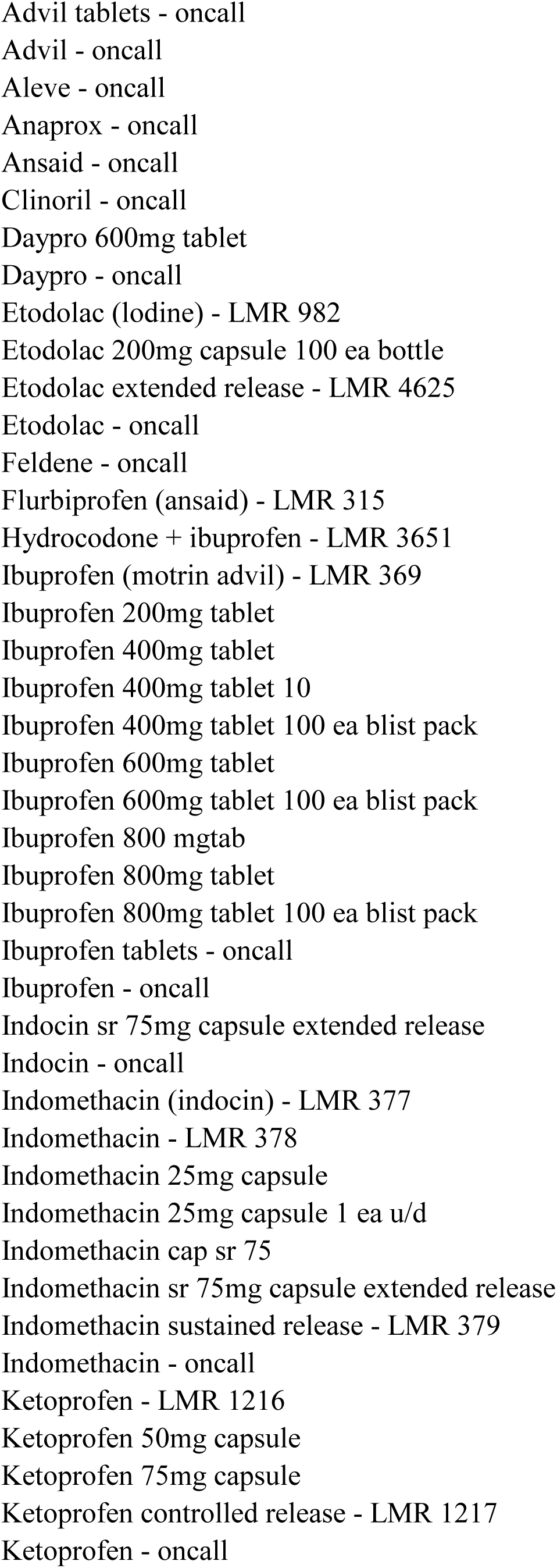
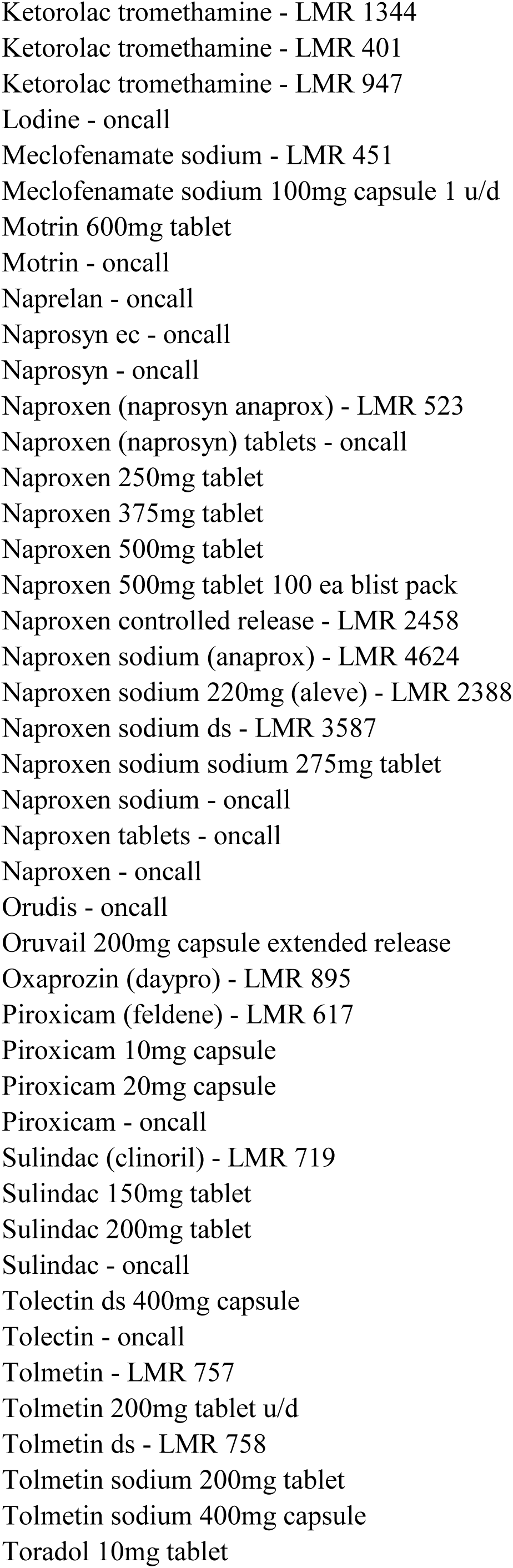
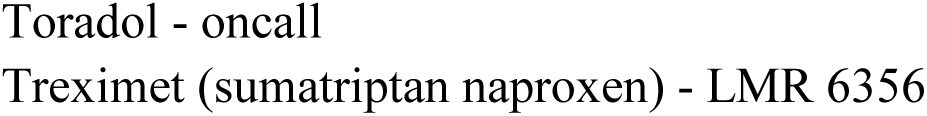
Nonselective NSAID prescriptions extracted from the EMR database

**Table S11.**
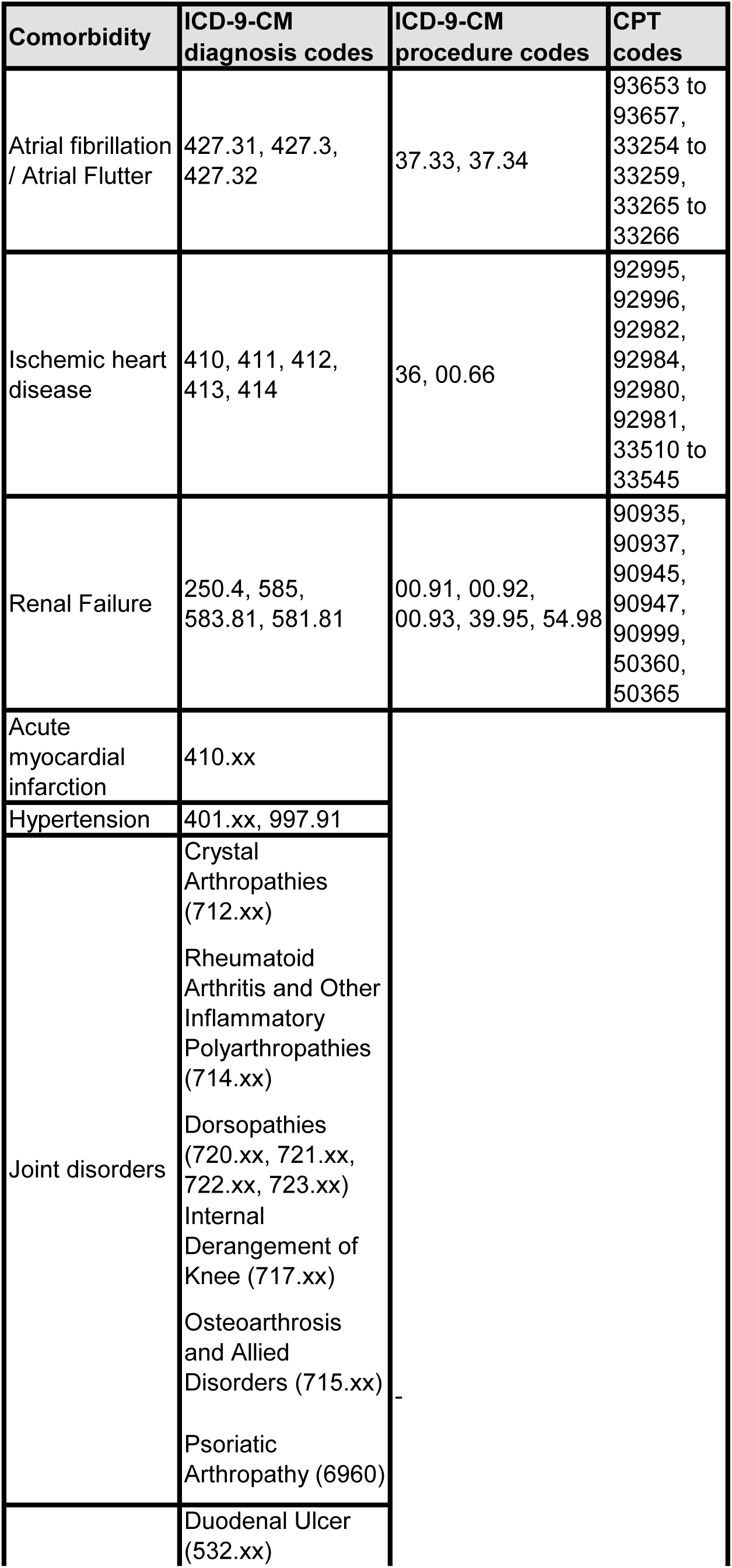
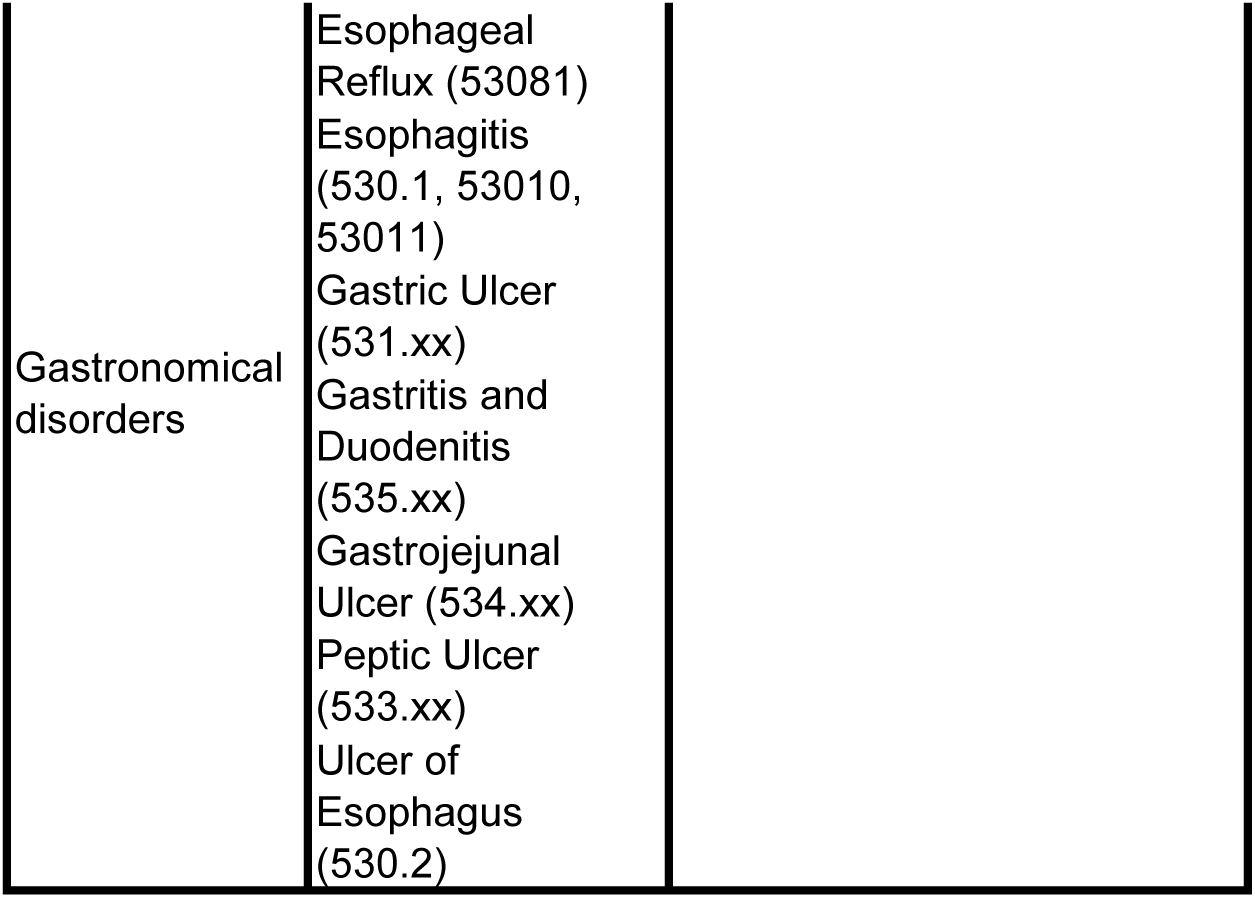
Diagnosis codes used to identify comorbidities in the EMR database.

**Table S12.**
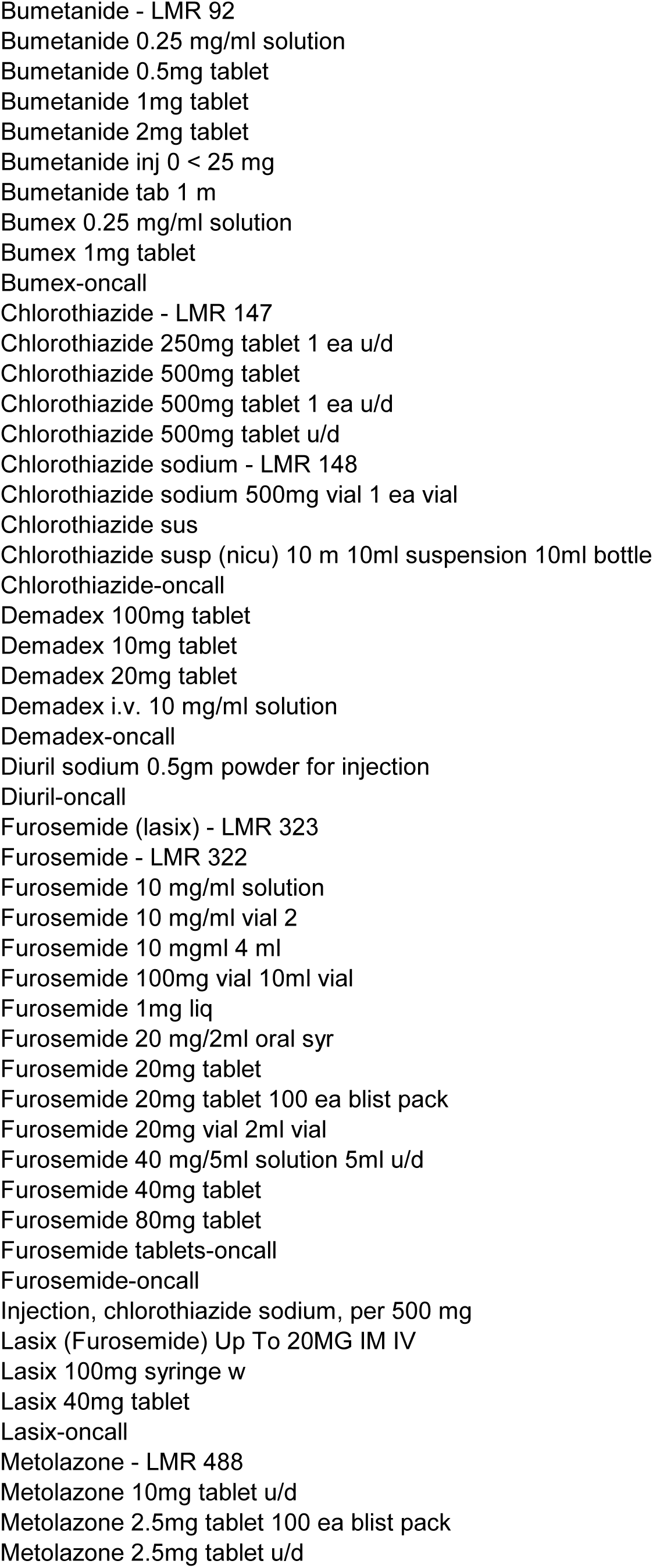
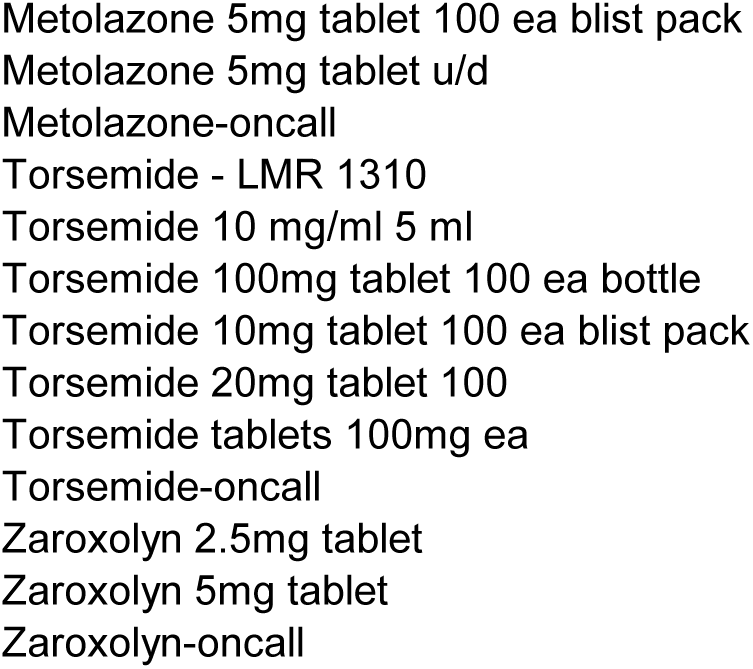
Diuretics prescriptions extracted from the EMR database

## Notes

### Competing Interest Statement

The authors have declared no competing interest.

### Summary of Updates

Text and figures revised. Supplemental tables updated.

